# mt-LAF3 is a pseudouridine synthase ortholog required for mitochondrial rRNA and mRNA gene expression in *Trypanosoma brucei*

**DOI:** 10.1101/2023.02.23.529727

**Authors:** Suzanne M McDermott, Vy Pham, Isaac Lewis, Maxwell Tracy, Kenneth Stuart

## Abstract

*Trypanosoma brucei* and related kinetoplastid parasites possess unique RNA processing pathways, including in their mitochondria, that regulate metabolism and development. Altering RNA composition or conformation through nucleotide modifications is one such pathway, and modifications including pseudouridine regulate RNA fate and function in many organisms. We surveyed pseudouridine synthase (PUS) orthologs in Trypanosomatids, with a particular interest in mitochondrial enzymes due to their potential importance for mitochondrial function and metabolism. *T. brucei* mt-LAF3 is an ortholog of human and yeast mitochondrial PUS enzymes, and a mitoribosome assembly factor, but structural studies differ in their conclusion as to whether it has PUS catalytic activity. Here, we generated *T. brucei* cells that are conditionally null for mt-LAF3 and showed that mt-LAF3 loss is lethal and disrupts mitochondrial membrane potential (ΔΨm). Addition of a mutant gamma-ATP synthase allele to the conditionally null cells permitted ΔΨm maintenance and cell survival, allowing us to assess primary effects on mitochondrial RNAs. As expected, these studies showed that loss of mt-LAF3 dramatically decreases levels of mitochondrial 12S and 9S rRNAs. Notably, we also observed decreases in mitochondrial mRNA levels, including differential effects on edited vs. pre-edited mRNAs, indicating that mt-LAF3 is required for mitochondrial rRNA and mRNA processing, including of edited transcripts. To assess the importance of PUS catalytic activity in mt-LAF3 we mutated a conserved aspartate that is necessary for catalysis in other PUS enzymes and showed it is not essential for cell growth, or maintenance of ΔΨm and mitochondrial RNA levels. Together, these results indicate that mt-LAF3 is required for normal expression of mitochondrial mRNAs in addition to rRNAs, but that PUS catalytic activity is not required for these functions. Instead, our work, combined with previous structural studies, suggests that *T. brucei* mt-LAF3 acts as a mitochondrial RNA-stabilizing scaffold.

## 1. Introduction

Pseudouridine (Ψ) was one of the first RNA modifications to be discovered (Cohn, 1960) and is thought to be one of the most abundant, found in all domains of life. Ψ is a seemingly simple modification derived from base-specific isomerization of uridine (U) and occurs either via a snoRNA-guided mechanism requiring H/ACA RNPs containing the Cbf5 or dyskerin enzyme in yeast and humans respectively, or snoRNA independent mechanisms catalyzed by pseudouridine synthases, also called PUS enzymes (Spenkuch et al., 2014; Li et al., 2016; Rintala-Dempsey and Kothe, 2017). The enzymes that catalyze pseudouridylation are classified into six families TruA, TruB (which includes Cbf5), TruD, RsuA, RluA, and PUS10. However, RsuA-type enzymes have so far only been described in bacteria while the PUS10 enzymes are present in only a few archaeal and eukaryotic organisms.

When incorporated, Ψ can alter the chemical and physical properties of RNA via its capacity to form an additional hydrogen bond compared to uridine, which can change RNA secondary structure via increased base stacking and altered base pairing (Arnez and Steitz, 1994). Ψ modifications have traditionally been studied in the context of non-coding RNAs, including tRNA, rRNA, and snRNAs. The TΨC arm of tRNAs contains a conserved Ψ at position 55 that suggests a fundamental role in translation, while rRNAs have numerous Ψ modifications near the peptidyltransferase center, the subunit interface of the ribosome and the decoding site, which are thought to impact the accuracy of translation and ribosomal assembly (Liang et al., 2009). snRNA pseudouridylation is essential for spliceosome biogenesis and activity and may be important for branch site recognition during splicing (Yu et al., 1998; Newby and Greenbaum, 2002; Wu et al., 2016). Furthermore, technologies for identification and mapping of Ψ have more recently revealed its presence in yeast, human, and other eukaryote mRNAs (Carlile et al., 2014; Li et al., 2015; Antonicka et al., 2017; Nakamoto et al., 2017). Replacement of U with Ψ in synthetic mRNAs increases translational efficiency (Kariko et al., 2008), although the roles of Ψs in mRNAs are not comprehensively understood.

Importantly, PUS enzymes are found in different cellular compartments, and some are found in multiple compartments (Rintala-Dempsey and Kothe, 2017), reflecting roles in all stages of processing of different types of RNA. For example, human mitochondrial granules that contain newly synthesized mitochondrial (mt) RNA also contain the PUS enzymes RPUSD4 and TRUB2 (RluA and TruB families respectively), as well as RPUSD3, which is an RPUSD4 paralog that lacks key catalytic residues (Borchardt et al., 2020). Depletion of each of these proteins results in decreased pseudouridylation of mt 16S rRNA and mRNAs encoding complex IV components COXI and COXIII, and defects in mitochondrial ribosome and OXPHOS complex assembly and translation (Arroyo et al., 2016; Antonicka et al., 2017; Zaganelli et al., 2017). Thus, there are clear essential roles for Ψ, PUS enzymes, and non-catalytic PUS enzyme paralogs in expression of the mitochondrial transcriptome.

Trypanosomatids including *Trypanosoma brucei* (*T. brucei*) are important parasites that are health and economic threats to millions of people globally. They have complex life cycles in mammals and insects that demand their ability to efficiently control both nuclear and mitochondrial gene expression and adapt their metabolism and physiology to different environments. Trypanosomes possess unique RNA processing pathways not found in other eukaryotes, including mitochondrial guide RNA (gRNA) directed U-insertion/deletion mRNA editing (Read et al., 2016; Aphasizheva et al., 2020), polycistronic transcription (El-Sayed et al., 2005; Clayton, 2016; Clayton, 2019), and trans-splicing (De Lange et al., 1984; Clayton, 2019), and these pathways are key regulators of gene expression, metabolism, and development. It is therefore likely that RNA modifications, such as Ψ, are important for both nuclear and mitochondrial gene expression in Trypanosomes. Indeed, Ψ has been mapped on *T. brucei* nuclear-encoded mRNAs (Rajan et al., 2021), rRNAs (Chikne et al., 2016), snRNAs (Rajan et al., 2019), snoRNAs (Rajan et al., 2022), and tRNAs (Rajan et al., 2022), and others including vault and 7SL RNAs (Rajan et al., 2022). Interestingly, many Ψ sites differ between mammalian bloodstream (BF) and insect procyclic (PF) form *T. brucei,* suggesting that these modifications are developmentally regulated. Furthermore, there is evidence that Ψ strengthens RNA–RNA and RNA–protein interactions within *T. brucei* ribosomes and spliceosomal snRNPs depending on temperature (Chikne et al., 2016; Rajan et al., 2019), and can influence mRNA stability via binding of regulatory proteins (Rajan et al., 2021), thus indicating that Ψ modifications are involved in life-cycle adaptation of *T. brucei* to different conditions in their mammalian hosts and insect vector.

Despite all this, our general knowledge of Trypanosomatid pseudouridine synthases, particularly in mitochondria, is limited. In *T. brucei,* the most studied enzyme is Cbf5 (Barth et al., 2005; Chikne et al., 2016; Rajan et al., 2019), which is an ortholog of Cbf5/dyskerin found in H/ACA RNPs. Silencing of Cbf5 in *T. brucei* leads to destabilization of H/ACA snoRNAs, including the spliced leader-associated RNA1 (SLA1), with reductions in levels of Ψ modification and maturation of snRNAs, spliced-leader RNA, and rRNA, and inhibition of trans-splicing (Barth et al., 2005). During the preparation of this manuscript several snoRNA-independent PUS orthologs were also identified (Maran et al., 2021; Rajan et al., 2021), and silencing a small number of these enzymes resulted in reductions in Ψ levels in specific rRNAs, tRNAs, and mRNAs (Rajan et al., 2021).

Here, we broadly surveyed PUS orthologs in Trypanosomatids, with a particular interest in mitochondrial enzymes due to their potential importance for mitochondrial RNA processing, including RNA editing, and metabolism. We identified *T. brucei* mitochondrial-Large ribosomal subunit Assembly Factor 3, mt-LAF3 (Tb927.9.3350; also known as PUS5), as an ortholog of the human mitochondrial RluA PUS paralogs RPUSD3 and RPUSD4 and of mitochondrial PUS5 in *S. cerevisiae.* Previous structural and RNAi knockdown studies indicated that *T. brucei* mt-LAF3 is a mitochondrial ribosome assembly factor that binds 12S rRNA and is required to maintain steady state levels of both 12S and 9S rRNAs in PF cells (Jaskolowski et al., 2020; Soufari et al., 2020). mt-LAF3 contains a conserved aspartate residue that is essential for catalysis in other PUS proteins (Del Campo et al., 2001). However, differences in the identities and positions of bound nucleotides relative to this catalytic aspartate in different structures have meant that it is unclear whether mt-LAF3 has PUS catalytic activity or not (Jaskolowski et al., 2020; Soufari et al., 2020; Rajan et al., 2021). We extended these previous studies by generating a BF *T. brucei* cell line that is conditionally null for mt-LAF3, thus allowing for a more complete knockdown of mt-LAF3 than in earlier RNAi experiments. In our conditional null cells, loss of mt-LAF3 resulted in loss of cell viability and ΔΨm, and large reductions in the steady state levels of both 12S and 9S mt rRNAs as expected. Critically, we also observed decreases in mt mRNA levels, including differential effects on several edited vs. pre-edited mRNAs, resulting in particularly severe reductions in edited COII, COIII, and A6 mRNAs. We modified our conditional null cell line by addition of a mutant gamma-ATP synthase allele that allows maintenance of ΔΨm following loss of the mitochondrial genome in BF *T. brucei* and used it to show that decreases in mt rRNA and edited mRNA levels are not secondary effects of changes in ΔΨm. The mutant gamma-ATP synthase allele completely rescued the cell viability phenotype, also indicating that the essential functions of mt-LAF3 in BF *T. brucei* are primarily in the mitochondrion. Finally, we analyzed the effects of mutation of the conserved catalytic aspartate in mt-LAF3 and showed that this residue and therefore PUS catalytic activity is not essential for cell growth, or maintenance of ΔΨm and mt rRNA or mRNA levels in BF cells. Together, these results show that mt-LAF3 is essential for normal expression of mt mRNAs in addition to mt 12S and 9S rRNAs in BF *T. brucei,* but that PUS catalytic activity is not required for either of these functions. Thus, our work combined with that of others, suggests that mt-LAF3 lacks PUS catalytic activity in BF *T. brucei,* and instead may act as a mitochondrial RNA stabilizing scaffold.

## 2. Material and Methods

### 2.1. Transfection and growth of cells in vitro

BF cells were grown in HMI-9 (18) with 10% FBS at 37 °C, 5% CO2. T ransfections of BF cell lines with the Amaxa Nucleofector (Lonza) were carried out as described (20), with the exception that Tb-BSF buffer (90 mM sodium phosphate buffer (Na_2_HPO_4_/NaH_2_PO_4_), 5 mM KCl, 0.15 mM CaCl_2_, 50 mM HEPES pH 7.2) was used for nucleofection instead of the Human T Cell Nucleofector Kit (21). Unless otherwise stated, concentrations of drugs used for BF cell selection and tet-regulated expression of transgenes are as follows: 2.5 μg/mL G418, 5 μg/mL hygromycin, 2.5 μg/mL phleomycin, 0.5 μg/mL tet, 0.1 μg/mL puromycin, 5 μg/mL blasticidin, 25 μg/mL ganciclovir.

### 2.2. Generation of BF conditional null cell line

Wild-type mt-LAF3 (Tb927.9.3350) lacking the stop codon and flanked in frame with attB Gateway recombination sites, was PCR amplified using primers described in Supplemental Table 1. BP Clonase II (Thermo Fisher Scientific) was used to transfer the PCR product into the Gateway entry vector pDONR221. LR Clonase II (Thermo Fisher Scientific) was then used to transfer the sequence into the Gateway destination vector pLEW100V5(BLE)GW-CtermTAP (Merritt and Stuart, 2013), which contains the phleomycin resistance (BLE) selectable marker and allows for tet-regulatable expression of C-terminal TAP-tagged mt-LAF3 in the ribosomal DNA spacer region. The resulting pLEW100v5(BLE)-mt-LAF3-CtermTAP plasmid was linearized with *Not*I, transfected into BF SM427 cells, and transgenic lines were selected by phleomycin resistance. Tet-regulated expression of mt-LAF3-TAP was confirmed by Western blotting with PAP antibody, and with an anti-Tubulin antibody as a loading control (Supplemental Table 2). Preparation of DNA constructs and transfections for endogenous allele knockouts using blasticidin (BSD)- and puromycin (PAC)-HSVTK drug cassettes in the regulatable BF cell line were generated and carried out as previously described (McDermott et al., 2015; McDermott et al., 2019). Correct insertion of knockout cassettes was assessed by PCR using primers described in Supplemental Table 1.

### 2.3. Generation of BF conditional null cell line containing the Mutant Gamma ATP synthase allele or exclusively expressed mt-LAF3 mutant allele

The drug selection cassettes in the CN cell line were excised using transient expression of Cre recombinase from pLEW100Cre_del_tetO (Addgene plasmid 24019; a gift from George Cross) and selection with ganciclovir (Invivogen) as previously described (McDermott et al., 2015; Carnes et al., 2017). Correct excision of the BSD- and PAC-HSVTK drug cassettes was determined via PCR (Supplemental Table 1) and drug sensitivity assays. For expression of the Mutant Gamma ATP synthase allele, the Cre-recombinase treated CN cells were transfected with a *Not*I-linearized pEnT6+ATPaseGammaWT+3UTR construct that allows the replacement of an endogenous wild-type ATPase gamma subunit allele with a mutant allele containing the L262P mutation (a gift from Matthew K. Gould and Achim Schnaufer) (Kelly et al., 2007; Dean et al., 2013; Carnes et al., 2017), and selected with blasticidin.

Constructs to generate CN cell lines containing wild-type or D211A alleles for exclusive expression were prepared via LR Clonase II-mediated transfer of the wild-type mt-LAF3 sequence into the destination vector pHD1344tub(PAC)GWCterm3V5, that allows for constitutive expression of C-terminal 3xV5 tagged protein in the βtubulin locus. The resulting pHD1344tub(PAC)-mt-LAF3-Cterm3V5 plasmid was then used as a template for site-directed mutagenesis of D211 (QuikChange II kit; Agilent) using forward and reverse primers listed in Supplemental Table 1. *NotI* digested plasmids containing the wild-type or D211A alleles were transfected into the Cre-recombinase treated CN cells. Transgenic lines were selected by puromycin resistance, construct insertion assessed by PCR (Supplemental Table 1), and constitutive expression of mt-LAF3-3xV5 confirmed by Western blotting with anti-V5 antibody, and with an anti-Tubulin antibody as a loading control (Supplemental Table 2).

### 2.4. ΔΨm measurement via flow cytometry

Samples of trypanosome cultures grown in the presence or absence of tet were incubated with 260 nM rhodamine 123 (Rh123) for 20 min at 37 °C. Cells were harvested by centrifugation at 1,300 × g for 10 min and washed one time with 25 mM HEPES pH 7.6, 120 mM KCl, 0.15 mM CaCl_2_, 10 mM K2HPO4/KH2PO4 pH 7.6, 2 mM EDTA, 5 mM MgCl2, and 6 mM glucose. Fluorescence caused by Rh123 uptake was measured using a LSR II flow cytometer with FACSDiva software (BD Biosciences). Baseline fluorescence was established for each sample by preincubating an aliquot of cells with 10 μM trifluorocarbonylcyanide phenylhydrazone (FCCP) before adding Rh123; the FCCP concentration was maintained throughout the wash and flow cytometer steps.

### 2.5. RNA isolation and RT-qPCR analysis

Total RNA was harvested using TRIzol and treated with TURBO DNase (Thermo Fisher Scientific) according to manufacturer’s instructions. RNA integrity was confirmed using an RNA nanochip on a BioAnalyzer (Agilent Technologies). 2 μg of total RNA was reverse transcribed using TaqMan Reverse Transcription Reagents and MultiScribe Reverse Transcriptase (Thermo Fisher Scientific), pre-amplified in multiplex Specific Target Amplification (STA) reactions using TaqMan PreAmp Master Mix (Thermo Fisher Scientific) and treated with Exonuclease I (NEB). The abundances of reference never-edited, pre-edited and edited transcript cDNAs were then analyzed by high-throughput real-time PCR on the BioMark HD system as previously described (McDermott et al., 2015). Primers are described in Supplemental Table 1. Calculations of RNA levels in samples following 48 hours tet withdrawal, relative to the presence of tet, were done using the 2 [-ΔΔC(T)] method (Livak and Schmittgen, 2001) using TERT as an internal reference. Technical duplicates of each cDNA sample were assayed for each target and internal reference per experiment and C(T) data averaged before performing the 2 [-ΔΔC(T)] calculation. Experiments were repeated using at least three biological replicates.

### 2.6. Protein homology detection and multiple sequence alignments

Sequence and structure-based homology searches were performed using HHPred (Zimmermann et al., 2018; Gabler et al., 2020) and Dali (Holm, 2020a, b, 2022). Multiple sequence alignments were performed with Geneious software (version 2019.1.1) using the Clustal Omega algorithm (version 1.2.3) with default parameters. Amino acid sequences were obtained from TriTrypDB (Aslett et al., 2010) and Uniprot (UniProt, 2022) and their accession/ID numbers are detailed in Supplementary Table 3.

### 2.7. Protein structure comparisons

The 12S rRNA-bound mt-LAF3 crystal structure (PDB 6YXY, chains K and F) was modelled onto *Escherichia coli* RluA (PDB 2I82, chains A and E) and *Homo sapiens* RPUSD4 (PDB 5UBA, chain A) crystal structures using the Matchmaker function in Chimera (Pettersen et al., 2004).

## 3. Results

### 3.1. *T. brucei* Tb927.9.3350 is a RluA family mitochondrial PUS protein

We found multiple orthologs of PUS enzymes in addition to Cbf5 in the *T. brucei* genome that together represent the major eukaryotic families TruA, TruB, TruD, RluA, and PUS10 (Supplementary Figure 1) (Maran et al., 2021). These relationships confirm a recent report published during the preparation of this manuscript that Tb927.7.1510, Tb927.6.2970, Tb927.9.3350, and Tb927.10.9050 are the *T. brucei* orthologs of *S. cerevisiae* PUS1, 3, 5, and 7 respectively (Rajan et al., 2021), and we also identify additional Trypanosomatid orthologs of these and other eukaryotic PUS enzymes (Supplementary Figure 1). Tb927.9.3350 is particularly intriguing due to its mitochondrial localization, shown via endogenous gene tagging in TrypTag (http://tryptag.org/?query=Tb927.9.3350) (Dean et al., 2017), detection in mitochondrial proteomic studies (Zikova et al., 2008; Panigrahi et al., 2009; Niemann et al., 2013), and isolation in tandem affinity purifications of multiple mitochondrial RNA editing proteins in the presence and absence of RNA (Aphasizheva et al., 2014). Using a combination of sequence and structural-based searches (Zimmermann et al., 2018; Gabler et al., 2020; Holm, 2020a, b, 2022), we detected homology (identity and similarity respectively) between Tb927.9.3350 and *T. cruzi* BCY84_12019 (78.9% and 93.6%), *Leishmania major* LMJFC_010007900 (70.1% and 86.9%), a *T. brucei* paralog Tb927.3.2130 (12.6% and 31.7%), human RPUSD4 (16.3% and 41.3%) and RPUSD3 (15.1% and 37.4%), and *S. cerevisiae* PUS5 (14.0% and 34.4%) (Figure 1, Supplementary Figures 1 and 2, Supplementary Table 3), which are all targeted or predicted to be targeted to mitochondria. *S. cerevisiae* PUS5 has only one known target, which is in mt 12S rRNA (Ansmant et al., 2000; Carlile et al., 2014). Consistent with the roles of its human and yeast orthologs, previous RNAi silencing of Tb927.9.3350, annotated as PUS5 (Rajan et al., 2021), in PF cells resulted in a ~60% decrease in the levels of mt 12S rRNA pseudouridylation at position 1018 as well as affecting modification of a small number of sites in nuclear-encoded mRNAs (Rajan et al., 2021). Tb927.9.3350 has also been annotated as mt-LAF3 in two other studies (Jaskolowski et al., 2020; Soufari et al., 2020), where it was characterized as an assembly factor for the mitoribosome large subunit required for binding and maturation of the 12S rRNA peptidyltransferase center (Jaskolowski et al., 2020; Soufari et al., 2020). Interestingly, these studies differed in their evaluation of whether mt-LAF3 plays a catalytic PUS role or is an rRNA scaffold in the mitoribosome based on sequence differences with other RluA-type enzymes in the catalytic and nearby sites (Figure 1), and conflicting observations of interactions between the catalytic core of the protein and a substrate uridine residue (Jaskolowski et al., 2020; Soufari et al., 2020). RNAi silencing of the gene in PF cells in (Jaskolowski et al., 2020) resulted in a strong growth defect and reductions in the steady state levels of both 12S and 9S rRNAs, supporting roles in mitoribosomal small subunit maturation or mit rRNA post-transcriptional processing, in addition to large subunit maturation (Jaskolowski et al., 2020). We will continue to refer to Tb927.9.3350 as mt-LAF3.

**Figure 1.**
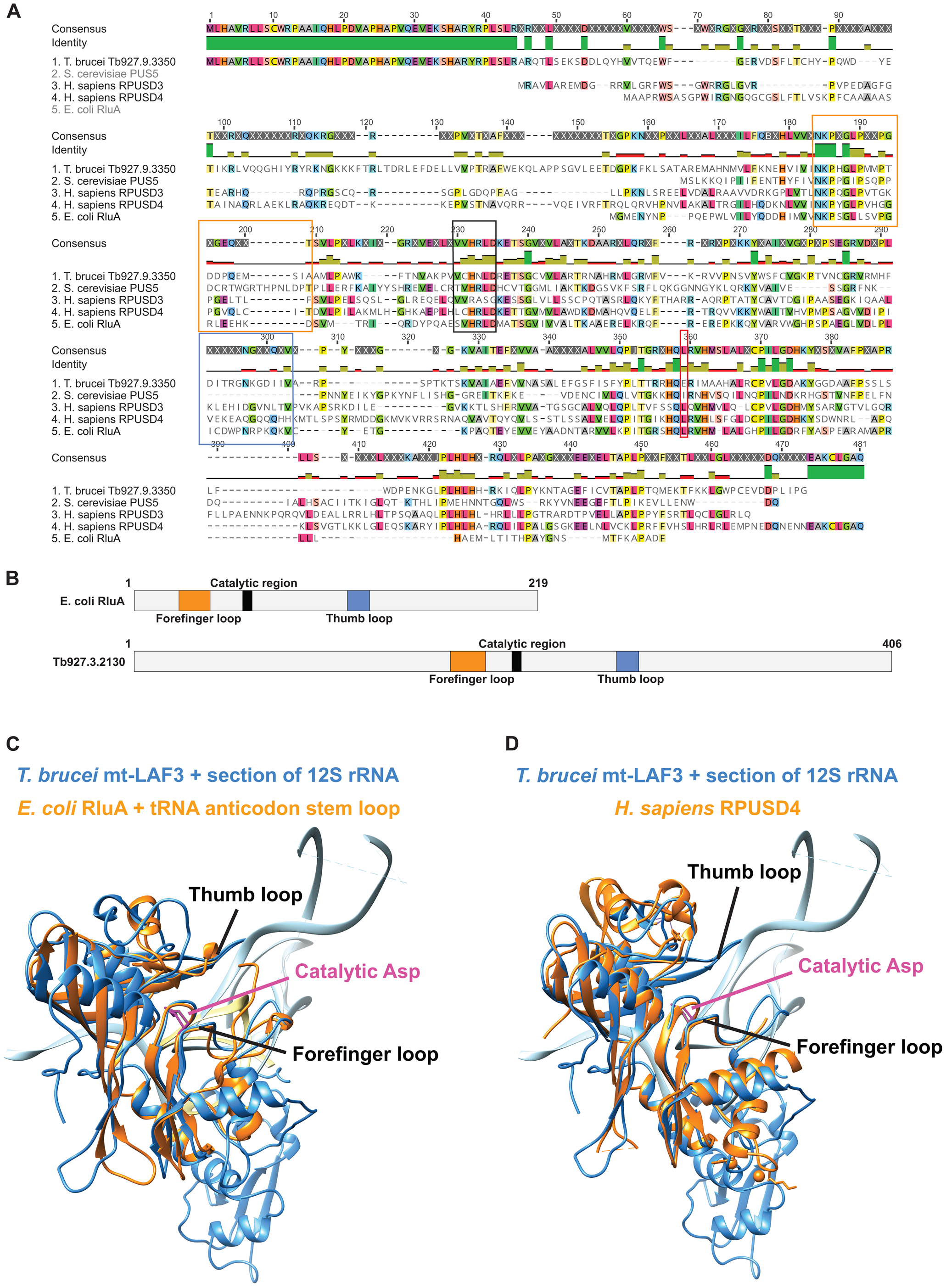
Tb927.9.3350/mt-LAF3 is related to RluA pseudouridine synthases. **(A)** Clustal Omega alignment of *T. brucei* Tb927.9.3350/mt-LAF3 with related representative orthologs from the RluA PUS family (Supplementary Table 3). The black box indicates the catalytic region, including the conserved catalytic aspartate (mt-LAF3 D211). An arginine residue thought to be key for RluA pseudouridine synthase activity (Hoang et al., 2006) is not conserved in mt-LAF3 (mt-LAF3 N209). Orange and blue boxes respectively surround the forefinger and thumb loops that pinch the RNA substrate in the RluA crystal structure (Hoang et al., 2006). The red box indicates an I/L residue found in most RluA family members that is not conserved in mt-LAF3 and its direct Trypanosomatid orthologs (mt-LAF3 E316). **(B)** Schematic of domain organization of *E. coli* RluA compared to *T. brucei* Tb927.9.3350/mt-LAF3 with key domains and motifs colored as in (A). **(C and D)** The crystal structure of *T. brucei* mt-LAF3 (PDB structure PDB 6YXY, chains K and F) shown in blue was compared to **(C)** the crystal structure of *E. coli* RluA (PDB 2I82, chains A and E) and **(D)** *H. sapiens* RPUSD4 (PDB 5UBA), both shown in orange. Bound RNAs are shown in light blue and orange respectively. Catalytic aspartate residues are highlighted in magenta, and the forefinger and thumb loops are also indicated.

### 3.2. mt-LAF3 is essential for cell growth and maintenance of mitochondrial membrane potential (△ψm) in BF *T. brucei*

We created a conditional null (CN) cell line in which both endogenous alleles of mt-LAF3 were deleted and in which a tetracycline (tet)-regulatable copy of mt-LAF3 was inserted into an ectopic ribosomal DNA locus. In the presence of tet, mt-LAF3 was expressed, and the cells grew in a similar manner to the parental BF 427 cell line. However, tet withdrawal led to a dramatic reduction in the expression of mt-LAF3 within one day and severe growth inhibition starting three days following removal of tet, indicating that mt-LAF3 is essential for BF growth (Figure 2A and Supplementary Figure 3). We also assayed mitochondrial function in response to mt-LAF3 knockdown via measurement of ΔΨm upon withdrawal of tet in both CN cells and CN cells containing the MGA allele. Live cells were stained with rhodamine 123 (Rh123), a fluorescent dye that is a marker for energized mitochondria (Divo et al., 1993; Wilkes et al., 1997), and the resultant green fluorescence intensity was measured using flow cytometry. Reduced fluorescence intensity and therefore ΔΨm corresponding to that seen upon treatment with the protonophore FCCP was observed in the CN cells two days following tet withdrawal and became more pronounced over time (Figure 2B and Supplementary Figure 4). Comparison of Rh123 fluorescence (and thus ΔΨm) with the growth curve in Figure 1C showed that the decrease of ΔΨm started at least 24 hours before the onset of the growth defect, suggesting that the decrease in ΔΨm is a primary response to mt-LAF3 loss and not a consequence of other lethal events.

**Figure 2.**
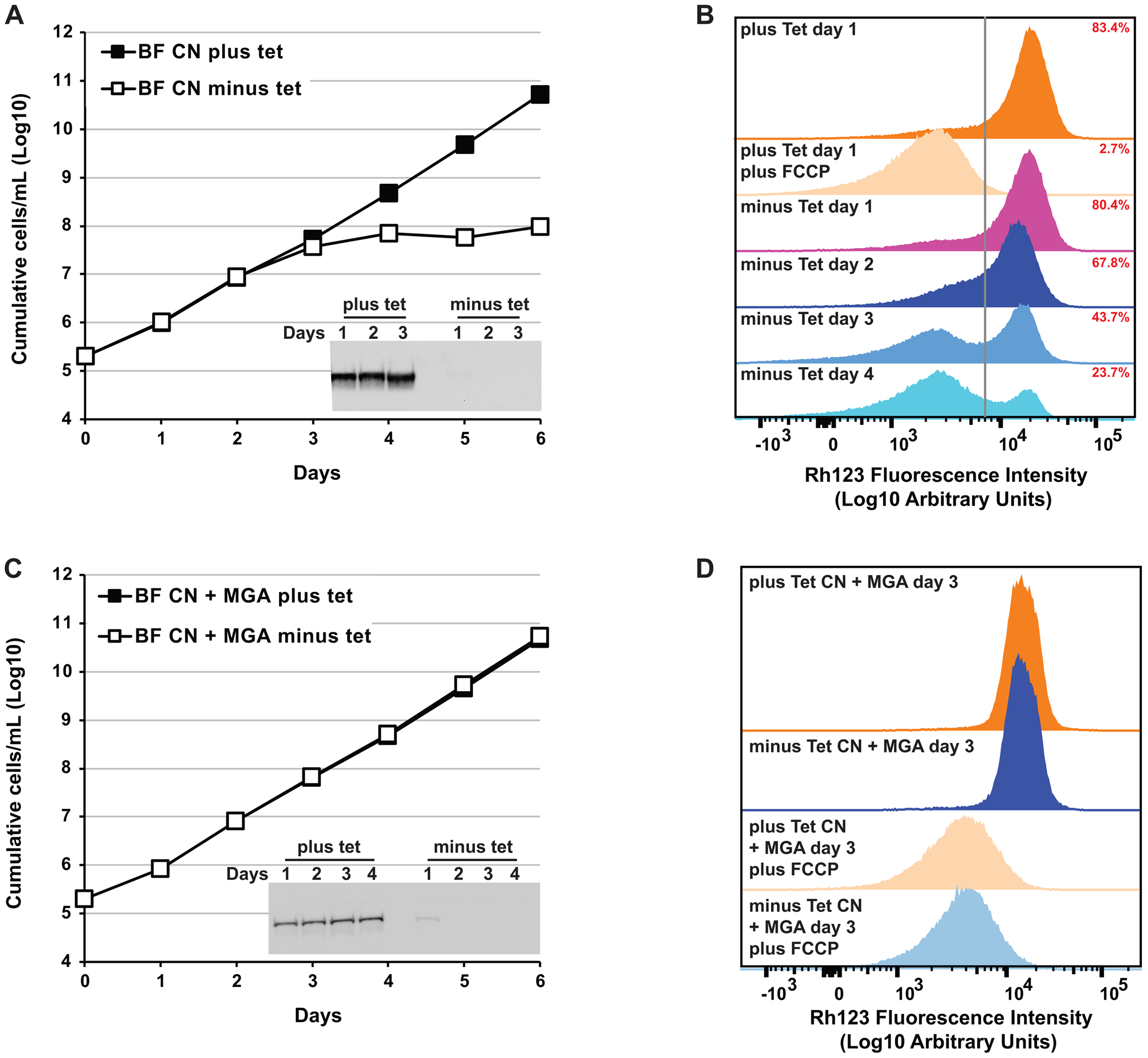
mt-LAF3 is essential for growth and maintenance of ΔΨm in BF *T. brucei* but the effects of mt-LAF3 loss can be circumvented by a L262P substitution in ATPase γ. BF mt-LAF3 CN cells that lack both endogenous mt-LAF3 alleles, and in which a tet-regulatable wild-type mt-LAF3 allele was expressed in the presence of tet or repressed in the absence of tet were generated. A mutant gamma-ATP synthase allele (L262P) that allows survival in the absence of mitochondrial gene expression was then transfected into the BF mt-LAF3 CN cells. **(A)** Cumulative growth of BF mt-LAF3 CN cells in the presence and absence of tet. Tet-regulated wild-type protein expression and repression were confirmed via Western blotting, probing with PAP antibody to allow detection of the TAP tag on the tet-regulatable wild-type mt-LAF3 (2 x 10^6^ cell equivalents per lane). **(B)** The effect of mt-LAF3 loss on the ΔΨm of the BF CN cells. ΔΨm was measured using rhodamine 123 (Rh123) and flow cytometry. The ‘ΔΨm negative’ gate (vertical line) was set to include counts detected for BF CN cells expressing mt-LAF3 in the presence of tet that were also pre-incubated with the protonophore trifluorocarbonylcyanide phenylhydrazone (FCCP). Counts to the right of this gate were defined as ‘ΔΨm positive’, and % cells determined to be ‘ΔΨm positive’ are shown in red. Representative analyses are shown, n = 2. **(C)** Cumulative growth of BF mt-LAF3 CN + MGA cells in the presence and absence of tet. Tet-regulated wild-type protein expression and repression were confirmed via Western blotting, probing with PAP antibody to allow detection of the TAP tag on the tet-regulatable wild-type mt-LAF3 (2 x 10^6^ cell equivalents per lane). **(D)** The effect of mt-LAF3 loss on the ΔΨm of the BF CN + MGA cells was measured using rhodamine 123 (Rh123) and flow cytometry as described for Figure 2B.

To assess whether the growth defect observed in these cells resulted solely from mitochondrial loss of function, we transfected a mutant gamma-ATP synthase allele (MGA; contains L262P substitution) into the mt-LAF3 CN cell line. This MGA allele was previously shown to compensate for the loss of the mitochondrial genome and gene expression in BF *T. brucei* (Dean et al., 2013; Carnes et al., 2017). The CN cells containing the MGA allele grew equally well in the presence and absence of tet meaning that the MGA allele was able to rescue growth in the absence of mt-LAF3 (Figure 2C). ΔΨm was also unchanged upon loss of mt-LAF3 in CN cells that contained the MGA allele (Figure 2D), indicating that the mutant gamma-ATP synthase was able participate in ΔΨm generation in cells that lack mt-LAF3 (Schnaufer et al., 2005; Dean et al., 2013). Together, these results indicate that the growth defect observed upon loss of mt-LAF3 in BF *T. brucei* is primarily due to loss of function in the mitochondrion. Furthermore, we show that loss of mt-LAF3 leads to a collapse of ΔΨm, which will impact essential mitochondrial functions (Zamzami et al., 1995a; Zamzami et al., 1995b; Schnaufer et al., 2002; Schnaufer et al., 2005; Kroemer et al., 2007; Moulin et al., 2019; Sato et al., 2019).

### 3.3. mt-LAF3 is required for normal mitochondrial rRNA and mRNA gene expression in BF *T. brucei*

Previous RNAi-mediated knockdown of mt-LAF3 in PF cells resulted in ~60% reductions in the steady state levels of both 12S and 9S mt rRNAs (Jaskolowski et al., 2020). A significant decrease in the levels of these transcripts and consequent effects on mitoribosomes and translation of ATP synthase could explain the growth and ΔΨm defects observed upon mt-LAF3 loss in BF. Here, we measured the steady state abundances of 12S and 9S rRNA transcripts by RT-qPCR in CN cells 36 hours following tet withdrawal relative to the same cells grown in the presence of tet. This timepoint was chosen to allow complete loss of mt-LAF3 mRNA and protein but precedes the ΔΨm and growth defects observed following tet withdrawal. The levels of 12S and 9S rRNAs were reduced much more dramatically than observed in previous studies, by ~95% and ~93% respectively (Figure 3). Importantly, mt-LAF3 mRNA was reduced by >99% confirming the loss of mt-LAF3 in our CN cell line.

**Figure 3.**
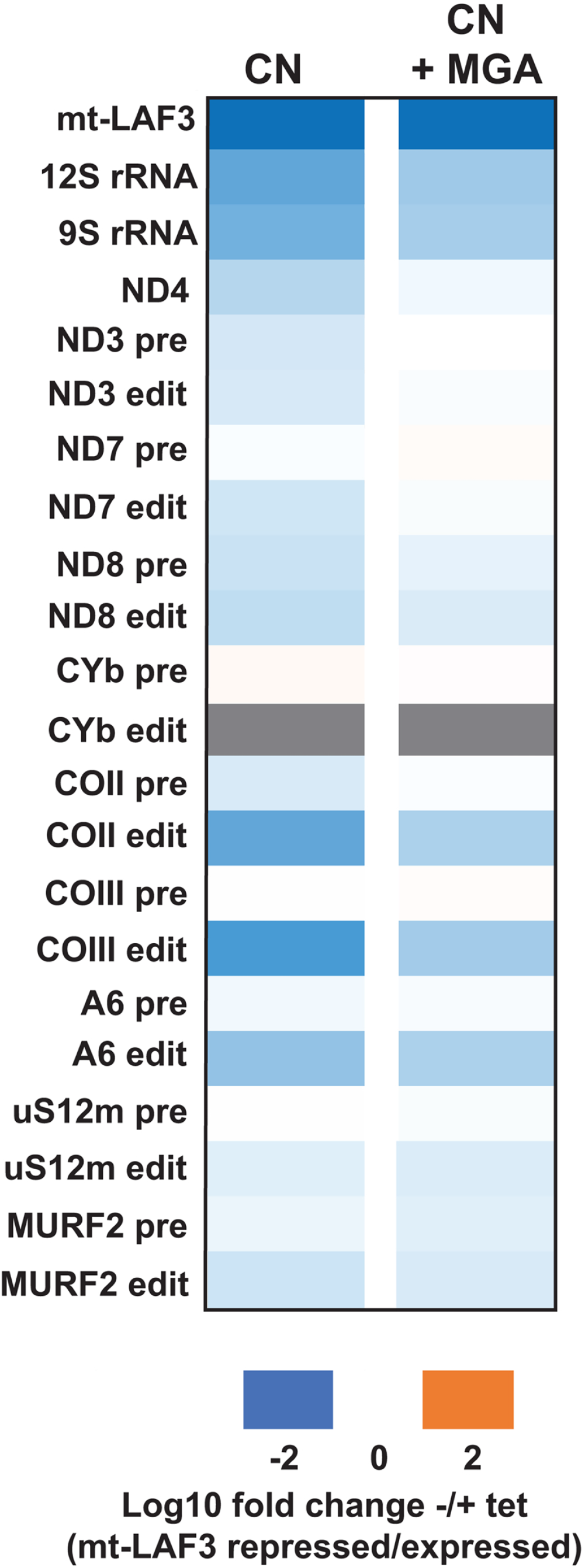
RT-qPCR analyses showing the impact of mt-LAF3 loss on mitochondrial RNAs, including pre-edited and edited mRNAs, *in vivo*. **(A)** RT-qPCR assay for a range of mitochondrial rRNAs and mRNAs in BF mt-LAF3 CN cells grown for 36 hours in the presence and absence of tet. For each target amplicon, the relative change in RNA abundance was determined by using telomerase reverse transcriptase (TERT) mRNA as an internal control, with BF CN cell lines that have either mt-LAF3 repressed compared to the same cell line in which mt-LAF3 was expressed. Gray box indicates that edited CYb was not detected in the CN cells grown either in the absence or presence of tet and we were therefore unable to quantify changes in its abundance. Data are shown as means from five independent experiments. **(B)** RT-qPCR assay as in (A) but examined in BF mt-LAF3 CN + MGA cells.

Given the roles of the human RPUSD3 and RPUSD4 orthologs in mt rRNA and mRNA expression, we hypothesized that mt-LAF3 may be involved in mt mRNA, in addition to rRNA, processing. To assess whether loss of mt-LAF3 impacted other mitochondrial transcripts in BF, we also measured the abundances of a range of other never-edited, pre-edited, and edited mRNAs by RT-qPCR in our CN cells 36 hours following tet withdrawal (Figure 3 and Supplementary Figure 5A). The level of never-edited ND4 was reduced by ~75%, and the abundances of pre-edited and edited ND3, ND7, ND8, and MURF2 were all reduced by ~60-70%. Interestingly, loss of mt-LAF3 had differential effects on several edited vs. pre-edited mRNAs, resulting in particularly severe reductions in edited COII, COIII, and A6 (~95%, ~96%, and ~86% respectively), with relatively small or moderate effects on the levels of pre-edited transcripts. We did not detect edited CYb mRNA in the BF cells grown either in the absence or presence of tet and so were unable to quantify changes in its abundance. We confirmed our qRT-PCR quantitation by amplifying and visualizing all pre-, partially-, and fully edited A6, ND3, ND8, and MURF2 RNAs in CN cells in the presence and absence of tet using RT-PCR and gel-electrophoresis. The results were consistent with RT-qPCR, as less fully edited, but more pre- and partially edited A6 were observed following loss of mt-LAF3 (Supplementary Figure 5B).

To address whether these changes in levels of mitochondrial rRNAs and mRNAs were primarily due to the loss of mt-LAF3 and not secondary consequences of changes in ΔΨm, we used the same RT-qPCR assays to analyze RNAs taken from the mt-LAF3 CN cells that also contain the MGA allele. Except for the mt-LAF3 transcript itself (>99% loss following tet withdrawal), the reductions in the steady state abundances of the assayed rRNA and mRNA transcripts were generally less severe than observed in the CN, indicating that the extent of RNA loss in the CN was in part due to the ΔΨm phenotype (Figure 3 and Supplementary Figure 5A). However, we again observed large reductions in the steady state abundances of 12S and 9S rRNAs upon following tet withdrawal (~86% and ~84% respectively), as well as in edited COII, COIII, and A6 (~78%, ~81%, and ~78% respectively) (Figure 3 and Supplementary Figure 5A). Thus, our results show that loss of mt-LAF3 in BF reduces the steady state levels of mitochondrial 12S and 9S rRNAs and mRNAs including edited COII, COIII, and A6 mRNAs, and is therefore required for mitochondrial rRNA and mRNA post-transcriptional processing.

### 3.4. Mutation of the conserved catalytic aspartate does not affect mt-LAF3 function in BF *T. brucei*

mt-LAF3 contains a conserved aspartate residue (D211) that is essential for catalysis in other PUS proteins (Del Campo et al., 2001). To address if the phenotypes we observed upon loss of mt-LAF3 were due to loss of PUS function, we asked whether expression of mt-LAF3 containing a D211A substitution could complement for the loss of mt-LAF3 or not. Wild-type, or mutant D211A mt-LAF3 alleles were tagged with a V5-epitope tag and inserted into the β-tubulin locus for constitutive expression in our BF CN cells. Western blotting confirmed that the V5-tagged alleles were expressed in the presence and absence of tet (Supplementary Figure 3). Exclusive expression of either the wild-type or D211A mutant allele was able to complement defects in growth, ΔΨm, and levels of mt rRNAs and mRNAs caused by loss of mt-LAF3 expression upon tet withdrawal from the CN cells (Figure 4). Thus, the conserved aspartate is not essential for normal mt rRNA or mRNA expression and BF cell growth, indicating that mt-LAF3 either lacks PUS activity or that its PUS activity is not required for these functions.

**Figure 4.**
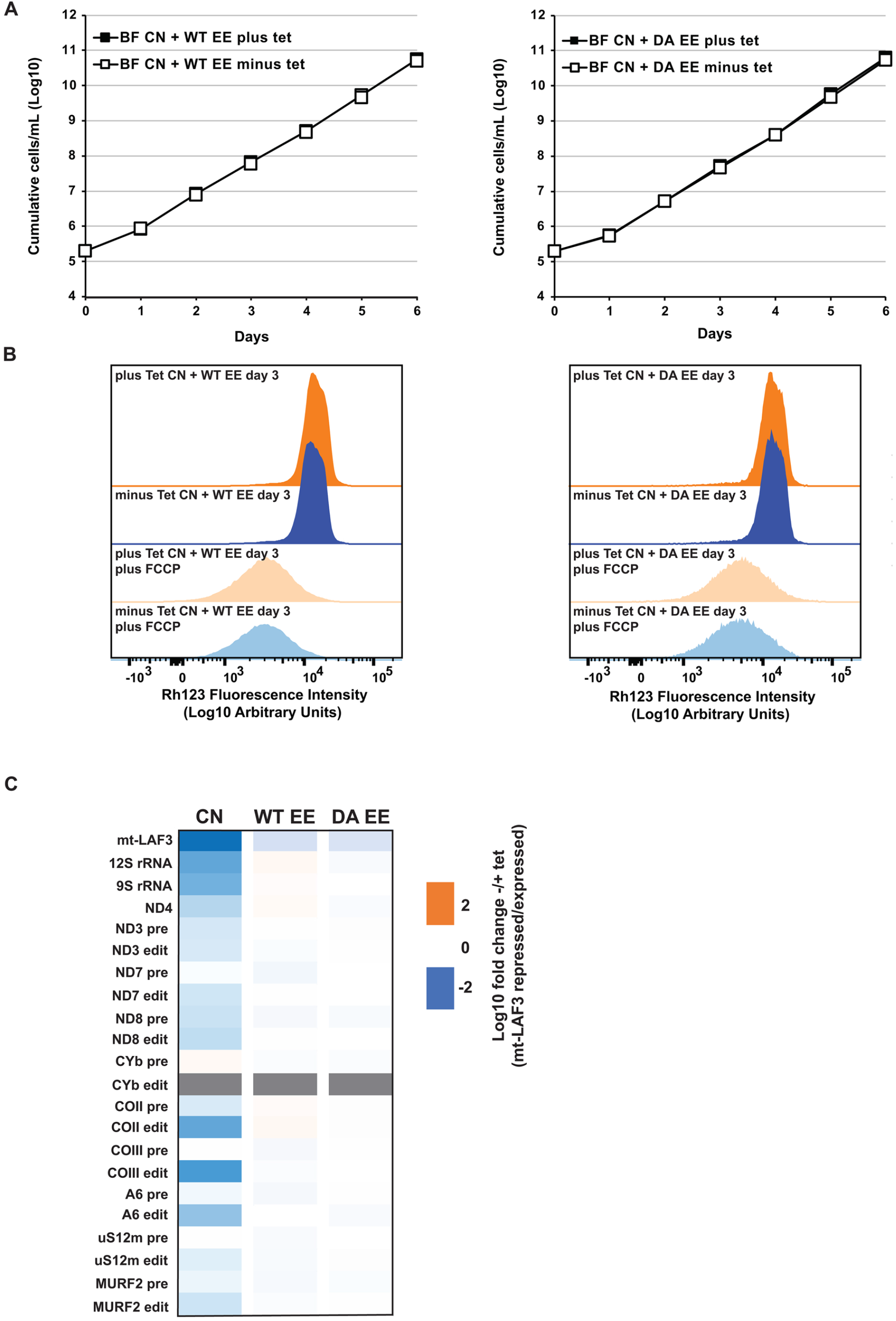
Mutation of the universally conserved catalytic aspartate in mt-LAF3 does not affect BF cell growth, ΔΨm, or abundances of mitochondrial RNAs. **(A)** Cumulative growth of BF mt-LAF3 CN cells that constitutively express V5-tagged wild-type (WT) or D211A (DA) mutant alleles, in the presence and absence of tet. In the absence of tet, the constitutive allele is exclusively expressed (EE). **(B)** The effect of three days of exclusive expression of WT or D211A (DA) mutant mt-LAF3 on the ΔΨm of the BF CN cells was measured using rhodamine 123 (Rh123) and flow cytometry as described in Figure 2B. **(C)** RT-qPCR assay for a range of mitochondrial rRNAs and mRNAs in BF mt-LAF3 CN cells (as shown in Figure 3A), and CN cells that exclusively express WT or D211A (DA) mutant mt-LAF3. Cells were grown for 36 hours in the presence and absence of tet, and their RNA levels measured relative to the telomerase reverse transcriptase (TERT) mRNA internal control. Gray box indicates that edited CYb was not detected in cells grown either in the absence or presence of tet and we were therefore unable to quantify changes in its abundance. WT and D211A exclusive expression data are shown as means from three independent experiments.

## 4. Discussion

Sequence and structure-based homology searches enabled us to identify multiple orthologs of PUS enzymes encoded in the *T. brucei* genome that together represent the major eukaryotic families TruA, TruB, TruD, RluA, and PUS10. One of these, Tb927.9.3350 or mt-LAF3, a mitochondrially localized protein also named PUS5 in previous studies, was particularly interesting given our focus on, and the limited information currently available for, mitochondrially targeted PUS enzymes in *T. brucei*. Furthermore, mt-LAF3 is an ortholog of the human mitochondrial RluA PUS paralogs RPUSD3 and RPUSD4, that are present in mtRNA granules where they are required for pseudouridylation of mt mRNA and rRNA respectively. Here, we made a conditional null BF *T. brucei* cell line and used it to show that loss of mt-LAF3 results in loss of cell viability and ΔΨm, and large reductions in the steady state levels of both 12S and 9S rRNAs. We expected that loss of mt-LAF3 would lead to reduced levels of mt 12S and 9S rRNAs given previous work showing the presence of mt-LAF3 in mitoribosomes (Jaskolowski et al., 2020; Soufari et al., 2020). However, we also observed reductions in the levels of edited COII, COIII, and A6 mRNAs that encode components of respiratory complexes III, IV, and V respectively. The edited forms, but not pre-edited forms of the mRNAs were specifically affected, indicating that mt-LAF3 is required for steps of mt mRNA, including of edited transcripts, in addition to rRNA post-transcriptional processing and maturation. Our results are consistent with a conserved role for mt-LAF3 in mt rRNA and mRNA gene expression, as depletion of the human orthologs RPUSD3 and RPUSD4 also resulted in defects in mitoribosome and OXPHOS complex component expression and assembly (Arroyo et al., 2016; Antonicka et al., 2017; Zaganelli et al., 2017).

Functions for mt-LAF3 in steps of mt mRNA, in addition to rRNA, processing are supported by its isolation in tandem affinity purifications of multiple mitochondrial RNA editing proteins, including RNA Editing Substrate Complex (RESC) components and KPAP1 poly(A) polymerase (Aphasizheva et al., 2014). Despite the somewhat different fates of mt rRNAs and edited mRNAs, it is possible that mt-LAF3 acts via similar mechanisms to impact their steady state levels in *T. brucei.* Edited mRNAs are modified by U-addition, with some U-deletion, progressively in a 3’ to 5’ direction in a process that requires the concerted actions of several large multiprotein complexes including RESCs and RNA Editing Core Complexes (RECCs) (Read et al., 2016; Aphasizheva et al., 2020). Edited messages are also marked by extension of initial short 20-30 nt A/U 3’-tails to long 200-300 nt A/U 3’-tails upon completion of editing by the combined activities of KRET1 TUTase and KPAP1 (Etheridge et al., 2008; Aphasizheva and Aphasizhev, 2010; Aphasizheva et al., 2011). The short tails apparently function in the regulation of transcript stability, while the long tails function at the interface of editing and translation but have not been shown to impact stability (Etheridge et al., 2008; Aphasizheva and Aphasizhev, 2010; Aphasizheva et al., 2011). Interestingly, both 12S and 9S rRNAs are also modified by KRET1-mediated U-addition at their 3’-end (Adler et al., 1991), and depletion of KRET1 results in a decline in their steady state levels (Aphasizheva and Aphasizhev, 2010). Thus mt-LAF3 may be part of the machinery that regulates the lengths and compositions of RNA tails and therefore stabilities in the *T. brucei* mitochondrion. The effects of 3’ tails on *T. brucei* mt mRNA stability are somewhat complex and can be transcript specific (Zimmer et al., 2012), which is consistent with our data that the steady state levels of only certain transcripts are affected upon loss of mt-LAF3. It is also possible that changes in edited mRNA levels are a secondary consequence of rRNA depletion and anticipated defects in ribosome assembly and translation. However, if this were the case, we might expect to see reductions in a greater range of edited transcripts, including those that are specifically fully edited in BF cells (Feagin and Stuart, 1985, 1988; Bhat et al., 1990; Koslowsky et al., 1990; Read et al., 1992; Souza et al., 1992; Souza et al., 1993; Corell et al., 1994; Read et al., 1994; Gazestani et al., 2018), and presumably translated. Furthermore, previous depletion of mitochondrial ribosome subunits did not generally affect mt mRNA levels, including of edited COII, and COIII, and A6 which were those transcripts specifically affected by loss of mt-LAF3 (Aphasizheva et al., 2014).

Substitution of the conserved catalytic aspartate D211 to alanine in *T. brucei* mt-LAF3 did not result in the growth, ΔΨm, or mtRNA abundance defects that we observed upon mt-LAF3 loss. This indicates that *T. brucei* mt-LAF3 either 1) lacks PUS activity or 2) that its PUS activity is not required for BF cell growth and normal mitochondrial gene expression in our assays. In most RluA PUS family members, the catalytic aspartate is found within a conserved HRLD motif. This motif contains both the catalytic aspartate and an arginine residue, which in RluA mediates the flipping of the target uridine into the catalytic pocket (Hoang et al., 2006). In mt-LAF3 this motif is HNLD, meaning that the catalytic aspartate is conserved, but that the uridine-flipping arginine residue required for PUS activity is not. Additional sequence differences between mt-LAF3 and other RluA PUS enzymes may also indicate lack of mt-LAF3 catalytic activity. These include a leucine or isoleucine residue in the vicinity of the RNA binding pocket in other RluA PUS proteins. mt-LAF3 lacks leucine or isoleucine in this position, and instead contains a glutamate residue (E316) that structurally points toward the active site and appears to hydrogen bond with the bound nucleotide, stabilizing binding of the nucleotide but tilting it away from the catalytic motif (Jaskolowski et al., 2020). Thus, these sequence and structural data are consistent with our mutational data, and together they suggest that mt-LAF3 lacks PUS catalytic activity, but instead acts as an rRNA- and mRNA-binding and stabilizing scaffold.

Lack of PUS activity in mt-LAF3 is not consistent with a previous study (Rajan et al., 2021) that reported decreases in pseudouridylation of mt 12S rRNA and some cytoplasmic mRNAs upon RNAi knockdown of mt-LAF3 in PF cells (Rajan et al., 2021). However, it is possible that there are life-cycle stage differences in mt-LAF3 catalytic function, as our study was conducted in BF cells, while the previous work was carried out in PF cells. A second possibility is that mt-LAF3 completely lacks PUS activity, but instead is required for interaction with, and recruitment of, active PUS enzymes, as has been described for its human orthologs RPUSD3 and RPUSD4. RPUSD3 and RPUSD4, together with a third TruB family PUS protein TRUB2, interact within human mtRNA granules (Antonicka et al., 2017), and depletion of each of the three individual proteins results in decreased pseudouridylation of mt rRNA or mRNAs. Interestingly, alignment of RPUSD3 and RPUSD4 with other RluA family members reveals that RPUSD3, like mt-LAF3, contains a divergent catalytic motif. RPUSD3 completely lacks HRLD, and instead contains the residues RASG. Notably, RPUSD3 has a glycine at the conserved position for the catalytic aspartate suggesting that RPUSD3 is not an active PUS enzyme (Borchardt et al., 2020), but instead is required for the catalytic activities of RPUSD4 and/or TRUB2. In our homology searches we also identified Tb927.3.2130 as a second predicted mitochondrial PUS in *T. brucei,* and a mt-LAF3 paralog. Tb927.3.2130 has all key residues found in active RluA PUS proteins, including the HRLD catalytic motif and an isoleucine in the vicinity of the RNA binding pocket, thus raising the possibility that mt-LAF3 and Tb927.3.2130 function analogously to human RPUSD3 and RPUSD4 respectively.

Overall, our work supports the idea that conserved mitochondrial PUS proteins and their non-catalytic paralogs are critical for mitochondrial RNA processing, including RNA editing, and mitochondrial function in *T. brucei.* It also raises the exciting possibility that there are more proteins involved in Trypanosomatid mitochondrial RNA processing to be identified despite detailed characterization of many key proteins and complexes (Gazestani et al., 2016; Aphasizheva et al., 2020).

## Acknowledgements

We thank Jason Carnes for helpful discussions and suggestions to improve the manuscript. This work was supported by NIH grant AI014102 to K.S.

## References

Adler, B.K., Harris, M.E., Bertrand, K.I., Hajduk, S.L., 1991. Modification of Trypanosoma brucei mitochondrial rRNA by posttranscriptional 3’ polyuridine tail formation. Mol Cell Biol 11, 5878–5884. doi: 10.1128/mcb.11.12.5878-5884.1991

Ansmant, I., Massenet, S., Grosjean, H., Motorin, Y., Branlant, C., 2000. Identification of the Saccharomyces cerevisiae RNA:pseudouridine synthase responsible for formation of psi(2819) in 21S mitochondrial ribosomal RNA. Nucleic Acids Res 28, 1941–1946. doi: 10.1093/nar/28.9.1941

Antonicka, H., Choquet, K., Lin, Z.Y., Gingras, A.C., Kleinman, C.L., Shoubridge, E.A., 2017. A pseudouridine synthase module is essential for mitochondrial protein synthesis and cell viability. EMBO Rep 18, 28–38. doi: 10.15252/embr.201643391

Aphasizheva, I., Alfonzo, J., Carnes, J., Cestari, I., Cruz-Reyes, J., Goringer, H.U., Hajduk, S., Lukes, J., Madison-Antenucci, S., Maslov, D.A., McDermott, S.M., Ochsenreiter, T., Read, L.K., Salavati, R., Schnaufer, A., Schneider, A., Simpson, L., Stuart, K., Yurchenko, V., Zhou, Z.H., Zikova, A., Zhang, L., Zimmer, S., Aphasizhev, R., 2020. Lexis and Grammar of Mitochondrial RNA Processing in Trypanosomes. Trends Parasitol 36, 337–355. doi: 10.1016/j.pt.2020.01.006

Aphasizheva, I., Aphasizhev, R., 2010. RET1-catalyzed uridylylation shapes the mitochondrial transcriptome in Trypanosoma brucei. Mol Cell Biol 30, 1555–1567. doi: 10.1128/MCB.01281-09

Aphasizheva, I., Maslov, D., Wang, X., Huang, L., Aphasizhev, R., 2011. Pentatricopeptide repeat proteins stimulate mRNA adenylation/uridylation to activate mitochondrial translation in trypanosomes. Mol Cell 42, 106–117. doi: 10.1016/j.molcel.2011.02.021

Aphasizheva, I., Zhang, L., Wang, X., Kaake, R.M., Huang, L., Monti, S., Aphasizhev, R., 2014. RNA binding and core complexes constitute the U-insertion/deletion editosome. Mol Cell Biol 34, 4329–4342. doi: 10.1128/MCB.01075-14

Arnez, J.G., Steitz, T.A., 1994. Crystal structure of unmodified tRNA(Gln) complexed with glutaminyl-tRNA synthetase and ATP suggests a possible role for pseudo-uridines in stabilization of RNA structure. Biochemistry 33, 7560–7567. doi: 10.1021/bi00190a008

Arroyo, J.D., Jourdain, A.A., Calvo, S.E., Ballarano, C.A., Doench, J.G., Root, D.E., Mootha, V.K., 2016. A Genome-wide CRISPR Death Screen Identifies Genes Essential for Oxidative Phosphorylation. Cell Metab 24, 875–885. doi: 10.1016/j.cmet.2016.08.017

Aslett, M., Aurrecoechea, C., Berriman, M., Brestelli, J., Brunk, B.P., Carrington, M., Depledge, D.P., Fischer, S., Gajria, B., Gao, X., Gardner, M.J., Gingle, A., Grant, G., Harb, O.S., Heiges, M., Hertz-Fowler, C., Houston, R., Innamorato, F., Iodice, J., Kissinger, J.C., Kraemer, E., Li, W., Logan, F.J., Miller, J.A., Mitra, S., Myler, P.J., Nayak, V., Pennington, C., Phan, I., Pinney, D.F., Ramasamy, G., Rogers, M.B., Roos, D.S., Ross, C., Sivam, D., Smith, D.F., Srinivasamoorthy, G., Stoeckert, C.J., Jr., Subramanian, S., Thibodeau, R., Tivey, A., Treatman, C., Velarde, G., Wang, H., 2010. TriTrypDB: a functional genomic resource for the Trypanosomatidae. Nucleic acids research 38, D457–462. doi: 10.1093/nar/gkp851

Barth, S., Hury, A., Liang, X.H., Michaeli, S., 2005. Elucidating the role of H/ACA-like RNAs in trans-splicing and rRNA processing via RNA interference silencing of the Trypanosoma brucei CBF5 pseudouridine synthase. J Biol Chem 280, 34558–34568. doi: 10.1074/jbc.M503465200

Bhat, G.J., Koslowsky, D.J., Feagin, J.E., Smiley, B.L., Stuart, K., 1990. An extensively edited mitochondrial transcript in kinetoplastids encodes a protein homologous to ATPase subunit 6. Cell 61, 885–894. doi: 10.1016/0092-8674(90)90199-o

Borchardt, E.K., Martinez, N.M., Gilbert, W.V., 2020. Regulation and Function of RNA Pseudouridylation in Human Cells. Annu Rev Genet 54, 309–336. doi: 10.1146/annurev-genet-112618-043830

Carlile, T.M., Rojas-Duran, M.F., Zinshteyn, B., Shin, H., Bartoli, K.M., Gilbert, W.V., 2014. Pseudouridine profiling reveals regulated mRNA pseudouridylation in yeast and human cells. Nature 515, 143–146. doi: 10.1038/nature13802

Carnes, J., McDermott, S., Anupama, A., Oliver, B.G., Sather, D.N., Stuart, K., 2017. In vivo cleavage specificity of Trypanosoma brucei editosome endonucleases. Nucleic Acids Res 45, 4667–4686. doi: 10.1093/nar/gkx116

Chikne, V., Doniger, T., Rajan, K.S., Bartok, O., Eliaz, D., Cohen-Chalamish, S., Tschudi, C., Unger, R., Hashem, Y., Kadener, S., Michaeli, S., 2016. A pseudouridylation switch in rRNA is implicated in ribosome function during the life cycle of Trypanosoma brucei. Sci Rep 6, 25296. doi: 10.1038/srep25296

Clayton, C., 2019. Regulation of gene expression in trypanosomatids: living with polycistronic transcription. Open Biol 9, 190072. doi: 10.1098/rsob.190072

Clayton, C.E., 2016. Gene expression in Kinetoplastids. Curr Opin Microbiol 32, 46–51. doi: 10.1016/j.mib.2016.04.018

Cohn, W.E., 1960. Pseudouridine, a carbon-carbon linked ribonucleoside in ribonucleic acids: isolation, structure, and chemical characteristics. J Biol Chem 235, 1488–1498. doi:

Corell, R.A., Myler, P., Stuart, K., 1994. Trypanosoma brucei mitochondrial CR4 gene encodes an extensively edited mRNA with completely edited sequence only in bloodstream forms. Mol Biochem Parasitol 64, 65–74. doi: 10.1016/0166-6851(94)90135-x

De Lange, T., Michels, P.A., Veerman, H.J., Cornelissen, A.W., Borst, P., 1984. Many trypanosome messenger RNAs share a common 5’ terminal sequence. Nucleic Acids Res 12, 3777–3790. doi: 10.1093/nar/12.9.3777

Dean, S., Gould, M.K., Dewar, C.E., Schnaufer, A.C., 2013. Single point mutations in ATP synthase compensate for mitochondrial genome loss in trypanosomes. Proc Natl Acad Sci U S A 110, 14741–14746. doi: 10.1073/pnas.1305404110

Dean, S., Sunter, J.D., Wheeler, R.J., 2017. TrypTag.org: A Trypanosome Genome-wide Protein Localisation Resource. Trends Parasitol 33, 80–82. doi: 10.1016/j.pt.2016.10.009

Del Campo, M., Kaya, Y., Ofengand, J., 2001. Identification and site of action of the remaining four putative pseudouridine synthases in Escherichia coli. RNA 7, 1603–1615. doi:

Divo, A.A., Patton, C.L., Sartorelli, A.C., 1993. Evaluation of rhodamine 123 as a probe for monitoring mitochondrial function in Trypanosoma brucei spp. J Eukaryot Microbiol 40, 329–335. doi: 10.1111/j.1550-7408.1993.tb04924.x

El-Sayed, N.M., Myler, P.J., Blandin, G., Berriman, M., Crabtree, J., Aggarwal, G., Caler, E., Renauld, H., Worthey, E.A., Hertz-Fowler, C., Ghedin, E., Peacock, C., Bartholomeu, D.C., Haas, B.J., Tran, A.N., Wortman, J.R., Alsmark, U.C., Angiuoli, S., Anupama, A., Badger, J., Bringaud, F., Cadag, E., Carlton, J.M., Cerqueira, G.C., Creasy, T., Delcher, A.L., Djikeng, A., Embley, T.M., Hauser, C., Ivens, A.C., Kummerfeld, S.K., Pereira-Leal, J.B., Nilsson, D., Peterson, J., Salzberg, S.L., Shallom, J., Silva, J.C., Sundaram, J., Westenberger, S., White, O., Melville, S.E., Donelson, J.E., Andersson, B., Stuart, K.D., Hall, N., 2005. Comparative genomics of trypanosomatid parasitic protozoa. Science 309, 404–409. doi: 10.1126/science.1112181

Etheridge, R.D., Aphasizheva, I., Gershon, P.D., Aphasizhev, R., 2008. 3’ adenylation determines mRNA abundance and monitors completion of RNA editing in T. brucei mitochondria. EMBO J 27, 1596–1608. doi: 10.1038/emboj.2008.87

Feagin, J.E., Stuart, K., 1985. Differential expression of mitochondrial genes between life cycle stages of Trypanosoma brucei. Proc Natl Acad Sci U S A 82, 3380–3384. doi: 10.1073/pnas.82.10.3380

Feagin, J.E., Stuart, K., 1988. Developmental aspects of uridine addition within mitochondrial transcripts of Trypanosoma brucei. Mol Cell Biol 8, 1259–1265. doi: 10.1128/mcb.8.3.1259-1265.1988

Gabler, F., Nam, S.Z., Till, S., Mirdita, M., Steinegger, M., Soding, J., Lupas, A.N., Alva, V., 2020. Protein Sequence Analysis Using the MPI Bioinformatics Toolkit. Curr Protoc Bioinformatics 72, e108. doi: 10.1002/cpbi.108

Gazestani, V.H., Hampton, M., Shaw, A.K., Salavati, R., Zimmer, S.L., 2018. Tail characteristics of Trypanosoma brucei mitochondrial transcripts are developmentally altered in a transcript-specific manner. Int J Parasitol 48, 179–189. doi: 10.1016/j.ijpara.2017.08.012

Gazestani, V.H., Nikpour, N., Mehta, V., Najafabadi, H.S., Moshiri, H., Jardim, A., Salavati, R., 2016. A Protein Complex Map of Trypanosoma brucei. PLoS Negl Trop Dis 10, e0004533. doi: 10.1371/journal.pntd.0004533

Hoang, C., Chen, J., Vizthum, C.A., Kandel, J.M., Hamilton, C.S., Mueller, E.G., Ferre-D’Amare, A.R., 2006. Crystal structure of pseudouridine synthase RluA: indirect sequence readout through protein-induced RNA structure. Mol Cell 24, 535–545. doi: 10.1016/j.molcel.2006.09.017

Holm, L., 2020a. DALI and the persistence of protein shape. Protein Sci 29, 128–140. doi: 10.1002/pro.3749

Holm, L., 2020b. Using Dali for Protein Structure Comparison. Methods Mol Biol 2112, 29–42. doi: 10.1007/978-1-0716-0270-6_3

Holm, L., 2022. Dali server: structural unification of protein families. Nucleic Acids Res 50, W210–215. doi: 10.1093/nar/gkac387

Jaskolowski, M., Ramrath, D.J.F., Bieri, P., Niemann, M., Mattei, S., Calderaro, S., Leibundgut, M., Horn, E.K., Boehringer, D., Schneider, A., Ban, N., 2020. Structural Insights into the Mechanism of Mitoribosomal Large Subunit Biogenesis. Mol Cell 79, 629–644 e624. doi: 10.1016/j.molcel.2020.06.030

Kariko, K., Muramatsu, H., Welsh, F.A., Ludwig, J., Kato, H., Akira, S., Weissman, D., 2008. Incorporation of pseudouridine into mRNA yields superior nonimmunogenic vector with increased translational capacity and biological stability. Mol Ther 16, 1833–1840. doi: 10.1038/mt.2008.200

Kelly, S., Reed, J., Kramer, S., Ellis, L., Webb, H., Sunter, J., Salje, J., Marinsek, N., Gull, K., Wickstead, B., Carrington, M., 2007. Functional genomics in Trypanosoma brucei: a collection of vectors for the expression of tagged proteins from endogenous and ectopic gene loci. Mol Biochem Parasitol 154, 103–109. doi: 10.1016/j.molbiopara.2007.03.012

Koslowsky, D.J., Bhat, G.J., Perrollaz, A.L., Feagin, J.E., Stuart, K., 1990. The MURF3 gene of T. brucei contains multiple domains of extensive editing and is homologous to a subunit of NADH dehydrogenase. Cell 62, 901–911. doi: 10.1016/0092-8674(90)90265-g

Kroemer, G., Galluzzi, L., Brenner, C., 2007. Mitochondrial membrane permeabilization in cell death. Physiol Rev 87, 99–163. doi: 10.1152/physrev.00013.2006

Li, X., Ma, S., Yi, C., 2016. Pseudouridine: the fifth RNA nucleotide with renewed interests. Curr Opin Chem Biol 33, 108–116. doi: 10.1016/j.cbpa.2016.06.014

Li, X., Zhu, P., Ma, S., Song, J., Bai, J., Sun, F., Yi, C., 2015. Chemical pulldown reveals dynamic pseudouridylation of the mammalian transcriptome. Nat Chem Biol 11, 592–597. doi: 10.1038/nchembio.1836

Liang, X.H., Liu, Q., Fournier, M.J., 2009. Loss of rRNA modifications in the decoding center of the ribosome impairs translation and strongly delays pre-rRNA processing. RNA 15, 1716–1728. doi: 10.1261/rna.1724409

Livak, K.J., Schmittgen, T.D., 2001. Analysis of relative gene expression data using real-time quantitative PCR and the 2(-Delta Delta C(T)) Method. Methods 25, 402–408. doi: 10.1006/meth.2001.1262

Maran, S.R., de Lemos Padilha Pitta, J.L., Dos Santos Vasconcelos, C.R., McDermott, S.M., Rezende, A.M., Silvio Moretti, N., 2021. Epitranscriptome machinery in Trypanosomatids: New players on the table? Mol Microbiol 115, 942–958. doi: 10.1111/mmi.14688

McDermott, S.M., Carnes, J., Stuart, K., 2019. Editosome RNase III domain interactions are essential for editing and differ between life cycle stages in Trypanosoma brucei. RNA 25, 1150–1163. doi: 10.1261/rna.071258.119

McDermott, S.M., Guo, X., Carnes, J., Stuart, K., 2015. Differential Editosome Protein Function between Life Cycle Stages of Trypanosoma brucei. J Biol Chem 290, 24914–24931. doi: 10.1074/jbc.M115.669432

Merritt, C., Stuart, K., 2013. Identification of essential and non-essential protein kinases by a fusion PCR method for efficient production of transgenic Trypanosoma brucei. Mol Biochem Parasitol 190, 44–49. doi: 10.1016/j.molbiopara.2013.05.002

Moulin, C., Caumont-Sarcos, A., Ieva, R., 2019. Mitochondrial presequence import: Multiple regulatory knobs fine-tune mitochondrial biogenesis and homeostasis. Biochim Biophys Acta Mol Cell Res 1866, 930–944. doi: 10.1016/j.bbamcr.2019.02.012

Nakamoto, M.A., Lovejoy, A.F., Cygan, A.M., Boothroyd, J.C., 2017. mRNA pseudouridylation affects RNA metabolism in the parasite Toxoplasma gondii. RNA 23, 1834–1849. doi: 10.1261/rna.062794.117

Newby, M.I., Greenbaum, N.L., 2002. Sculpting of the spliceosomal branch site recognition motif by a conserved pseudouridine. Nat Struct Biol 9, 958–965. doi: 10.1038/nsb873

Niemann, M., Wiese, S., Mani, J., Chanfon, A., Jackson, C., Meisinger, C., Warscheid, B., Schneider, A., 2013. Mitochondrial outer membrane proteome of Trypanosoma brucei reveals novel factors required to maintain mitochondrial morphology. Mol Cell Proteomics 12, 515–528. doi: 10.1074/mcp.M112.023093

Panigrahi, A.K., Ogata, Y., Zikova, A., Anupama, A., Dalley, R.A., Acestor, N., Myler, P.J., Stuart, K.D., 2009. A comprehensive analysis of Trypanosoma brucei mitochondrial proteome. Proteomics 9, 434–450. doi: 10.1002/pmic.200800477

Pettersen, E.F., Goddard, T.D., Huang, C.C., Couch, G.S., Greenblatt, D.M., Meng, E.C., Ferrin, T.E., 2004. UCSF Chimera--a visualization system for exploratory research and analysis. J Comput Chem 25, 1605–1612. doi: 10.1002/jcc.20084

Rajan, K.S., Adler, K., Doniger, T., Cohen-Chalamish, S., Aharon-Hefetz, N., Aryal, S., Pilpel, Y., Tschudi, C., Unger, R., Michaeli, S., 2022. Identification and functional implications of pseudouridine RNA modification on small noncoding RNAs in the mammalian pathogen Trypanosoma brucei. J Biol Chem 298, 102141. doi: 10.1016/j.jbc.2022.102141

Rajan, K.S., Adler, K., Madmoni, H., Peleg-Chen, D., Cohen-Chalamish, S., Doniger, T., Galili, B., Gerber, D., Unger, R., Tschudi, C., Michaeli, S., 2021. Pseudouridines on Trypanosoma brucei mRNAs are developmentally regulated: Implications to mRNA stability and protein binding. Mol Microbiol 116, 808–826. doi: 10.1111/mmi.14774

Rajan, K.S., Doniger, T., Cohen-Chalamish, S., Chen, D., Semo, O., Aryal, S., Glick Saar, E., Chikne, V., Gerber, D., Unger, R., Tschudi, C., Michaeli, S., 2019. Pseudouridines on Trypanosoma brucei spliceosomal small nuclear RNAs and their implication for RNA and protein interactions. Nucleic Acids Res 47, 7633–7647. doi: 10.1093/nar/gkz477

Read, L.K., Lukes, J., Hashimi, H., 2016. Trypanosome RNA editing: the complexity of getting U in and taking U out. Wiley Interdiscip Rev RNA 7, 33–51. doi: 10.1002/wrna.1313

Read, L.K., Myler, P.J., Stuart, K., 1992. Extensive editing of both processed and preprocessed maxicircle CR6 transcripts in Trypanosoma brucei. J Biol Chem 267, 1123–1128. doi:

Read, L.K., Wilson, K.D., Myler, P.J., Stuart, K., 1994. Editing of Trypanosoma brucei maxicircle CR5 mRNA generates variable carboxy terminal predicted protein sequences. Nucleic Acids Res 22, 1489–1495. doi: 10.1093/nar/22.8.1489

Rintala-Dempsey, A.C., Kothe, U., 2017. Eukaryotic stand-alone pseudouridine synthases - RNA modifying enzymes and emerging regulators of gene expression? RNA Biol 14, 1185–1196. doi: 10.1080/15476286.2016.1276150

Sato, T.K., Kawano, S., Endo, T., 2019. Role of the membrane potential in mitochondrial protein unfolding and import. Sci Rep 9, 7637. doi: 10.1038/s41598-019-44152-z

Schnaufer, A., Clark-Walker, G.D., Steinberg, A.G., Stuart, K., 2005. The F1-ATP synthase complex in bloodstream stage trypanosomes has an unusual and essential function. EMBO J 24, 4029–4040. doi: 10.1038/sj.emboj.7600862

Schnaufer, A., Domingo, G.J., Stuart, K., 2002. Natural and induced dyskinetoplastic trypanosomatids: how to live without mitochondrial DNA. Int J Parasitol 32, 1071–1084. doi: 10.1016/s0020-7519(02)00020-6

Soufari, H., Waltz, F., Parrot, C., Durrieu-Gaillard, S., Bochler, A., Kuhn, L., Sissler, M., Hashem, Y., 2020. Structure of the mature kinetoplastids mitoribosome and insights into its large subunit biogenesis. Proc Natl Acad Sci U S A 117, 29851–29861. doi: 10.1073/pnas.2011301117

Souza, A.E., Myler, P.J., Stuart, K., 1992. Maxicircle CR1 transcripts of Trypanosoma brucei are edited and developmentally regulated and encode a putative iron-sulfur protein homologous to an NADH dehydrogenase subunit. Mol Cell Biol 12, 2100–2107. doi: 10.1128/mcb.12.5.2100-2107.1992

Souza, A.E., Shu, H.H., Read, L.K., Myler, P.J., Stuart, K.D., 1993. Extensive editing of CR2 maxicircle transcripts of Trypanosoma brucei predicts a protein with homology to a subunit of NADH dehydrogenase. Mol Cell Biol 13, 6832–6840. doi: 10.1128/mcb.13.11.6832-6840.1993

Spenkuch, F., Motorin, Y., Helm, M., 2014. Pseudouridine: still mysterious, but never a fake (uridine)! RNA Biol 11, 1540–1554. doi: 10.4161/15476286.2014.992278

UniProt, C., 2022. UniProt: the Universal Protein Knowledgebase in 2023. Nucleic Acids Res. doi: 10.1093/nar/gkac1052

Wilkes, J.M., Mulugeta, W., Wells, C., Peregrine, A.S., 1997. Modulation of mitochondrial electrical potential: a candidate mechanism for drug resistance in African trypanosomes. Biochem J 326 (Pt 3), 755–761. doi: 10.1042/bj3260755

Wu, G., Adachi, H., Ge, J., Stephenson, D., Query, C.C., Yu, Y.T., 2016. Pseudouridines in U2 snRNA stimulate the ATPase activity of Prp5 during spliceosome assembly. EMBO J 35, 654–667. doi: 10.15252/embj.201593113

Yu, Y.T., Shu, M.D., Steitz, J.A., 1998. Modifications of U2 snRNA are required for snRNP assembly and pre-mRNA splicing. EMBO J 17, 5783–5795. doi: 10.1093/emboj/17.19.5783

Zaganelli, S., Rebelo-Guiomar, P., Maundrell, K., Rozanska, A., Pierredon, S., Powell, C.A., Jourdain, A.A., Hulo, N., Lightowlers, R.N., Chrzanowska-Lightowlers, Z.M., Minczuk, M., Martinou, J.C., 2017. The Pseudouridine Synthase RPUSD4 Is an Essential Component of Mitochondrial RNA Granules. J Biol Chem 292, 4519–4532. doi: 10.1074/jbc.M116.771105

Zamzami, N., Marchetti, P., Castedo, M., Decaudin, D., Macho, A., Hirsch, T., Susin, S.A., Petit, P.X., Mignotte, B., Kroemer, G., 1995a. Sequential reduction of mitochondrial transmembrane potential and generation of reactive oxygen species in early programmed cell death. J Exp Med 182, 367–377. doi: 10.1084/jem.182.2.367

Zamzami, N., Marchetti, P., Castedo, M., Zanin, C., Vayssiere, J.L., Petit, P.X., Kroemer, G., 1995b. Reduction in mitochondrial potential constitutes an early irreversible step of programmed lymphocyte death in vivo. J Exp Med 181, 1661–1672. doi: 10.1084/jem.181.5.1661

Zikova, A., Panigrahi, A.K., Dalley, R.A., Acestor, N., Anupama, A., Ogata, Y., Myler, P.J., Stuart, K., 2008. Trypanosoma brucei mitochondrial ribosomes: affinity purification and component identification by mass spectrometry. Mol Cell Proteomics 7, 1286–1296. doi: 10.1074/mcp.M700490-MCP200

Zimmer, S.L., McEvoy, S.M., Menon, S., Read, L.K., 2012. Additive and transcript-specific effects of KPAP1 and TbRND activities on 3’ non-encoded tail characteristics and mRNA stability in Trypanosoma brucei. PLoS One 7, e37639. doi: 10.1371/journal.pone.0037639

Zimmermann, L., Stephens, A., Nam, S.Z., Rau, D., Kubler, J., Lozajic, M., Gabler, F., Soding, J., Lupas, A.N., Alva, V., 2018. A Completely Reimplemented MPI Bioinformatics Toolkit with a New HHpred Server at its Core. J Mol Biol 430, 2237–2243. doi: 10.1016/j.jmb.2017.12.007

