## Supplementary Figures for "mt-LAF3 is a pseudouridine synthase ortholog required for mitochondrial rRNA and mRNA gene expression in *Trypanosoma brucei*"

**Supplementary Figure 1**

**A**

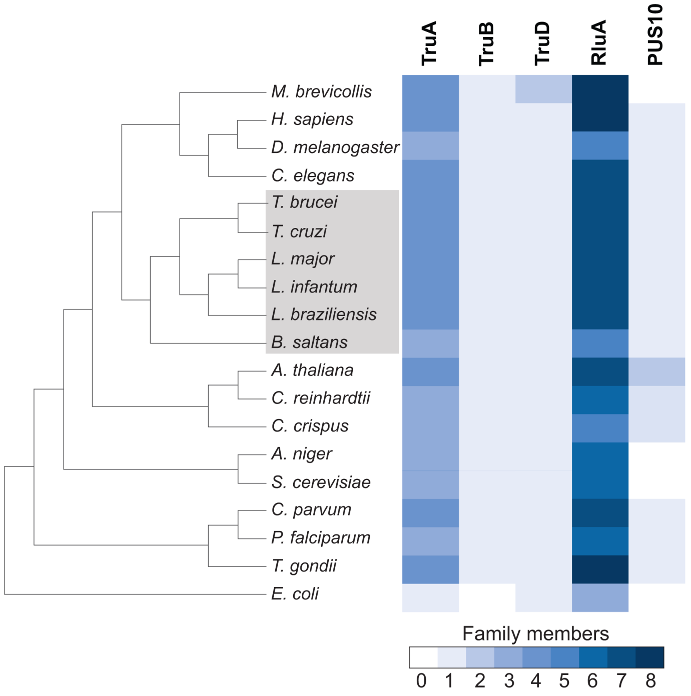


**B**

| % identity/ similarity | *T. brucei* mt-LAF3 Tb927.9.3350 | *T. cruzi* BCY84_12019 | *L. major* LMJFC_010007900 | *T. brucei* Tb927.3.2130 | *H. sapiens* RPUSD4 | *H. sapiens* RPUSD3 | *S. cerevisiae* PUS5 | *E. coli* RluA |
|---|---|---|---|---|---|---|---|---|
| *T. brucei* mt-LAF3 Tb927.9.3350 | | 78.9%/ 93.6% | 70.1%/ 86.9% | 12.6%/ 31.7% | 16.3%/ 41.3% | 15.1%/ 37.4% | 14.0%/ 34.4% | 21.3%/ 42.3% |
| *T. cruzi* BCY84_12019 | 78.9%/ 93.6% | | 72.9%/ 86.2% | 11.6%/ 31.1% | 16.3%/ 41.3% | 15.1%/ 37.8% | 13.0%/ 35.1% | 20.2%/ 42.3% |
| *L. major* LMJFC_010007900 | 70.1%/ 86.9% | 72.9%/ 86.2% | | 11.2%/ 29.7% | 16.8%/ 43.7% | 16.1%/ 38.8% | 13.4%/ 34.1% | 19.4%/ 41.1% |
| *T. brucei* Tb927.3.2130 | 12.6%/ 31.7% | 11.6%/ 31.1% | 11.2%/ 29.7% | | 16.8%/ 35.0% | 14.3%/ 33.1% | 13.4%/ 35.4% | 20.9%/ 42.2% |
| *H. sapiens* RPUSD4 | 16.3%/ 41.3% | 16.3%/ 41.3% | 16.8%/ 43.7% | 16.8%/ 35.0% | | 23.9%/ 50.1% | 16.6%/ 40.2% | 26.5%/ 49.3% |
| *H. sapiens* RPUSD3 | 15.1%/ 37.4% | 15.1%/ 37.8% | 16.1%/ 38.8% | 14.3%/ 33.1% | 23.9%/ 50.1% | | 10.1%/ 33.2% | 18.0%/ 42.1% |
| *S. cerevisiae* PUS5 | 14.0%/ 34.4% | 13.0%/ 35.1% | 13.4%/ 34.1% | 13.4%/ 35.4% | 16.6%/ 40.2% | 10.1%/ 33.2% | | 17.0%/ 38.0% |
| *E. coli* RluA | 21.3%/ 42.3% | 20.2%/ 42.3% | 19.4%/ 41.1% | 20.9%/ 42.2% | 26.5%/ 49.3% | 18.0%/ 42.1% | 17.0%/ 38.0% | |

### Supplementary Figure 2

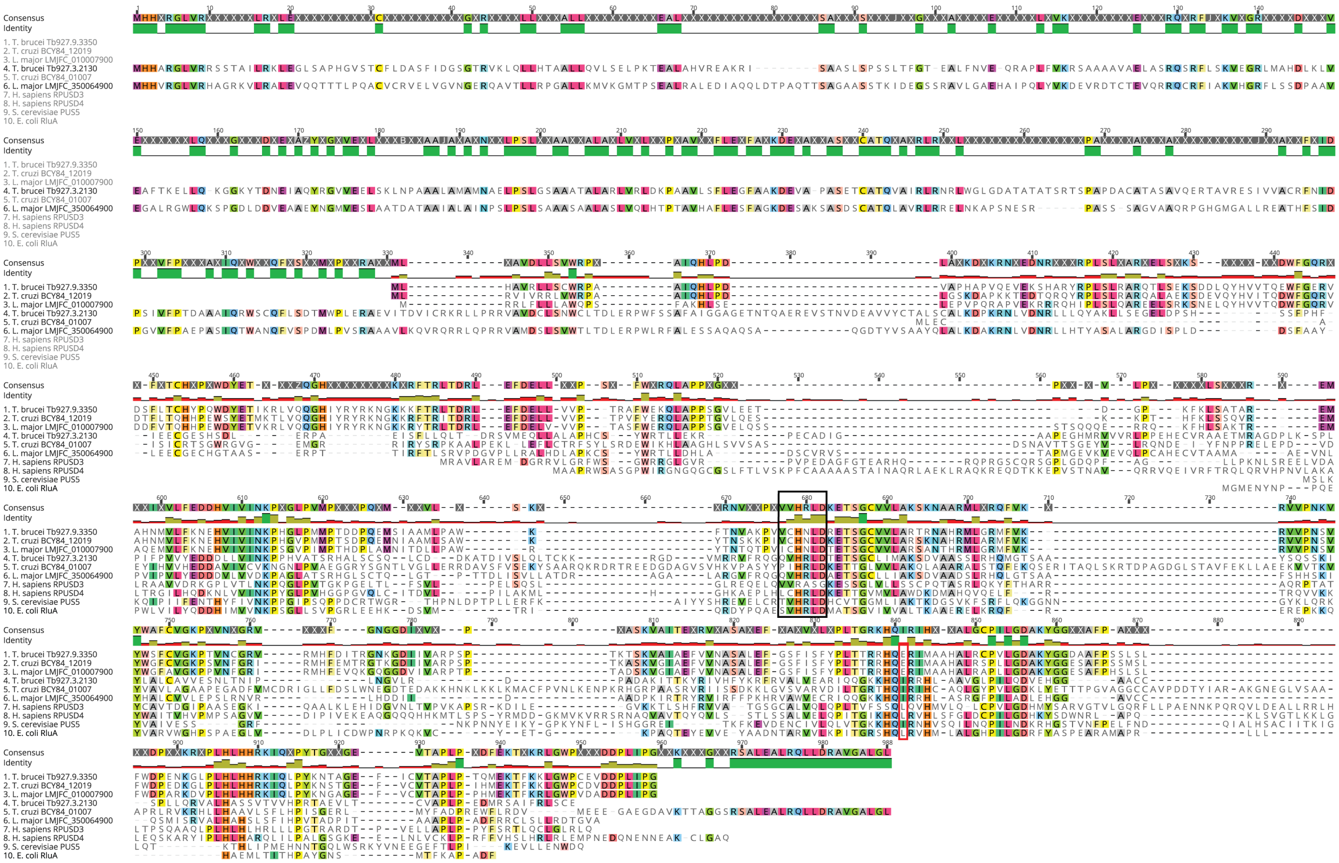


### Supplementary Figure 3

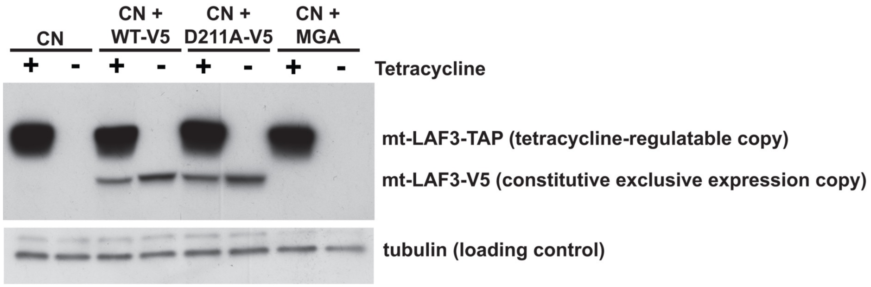

### Supplementary Figure 4

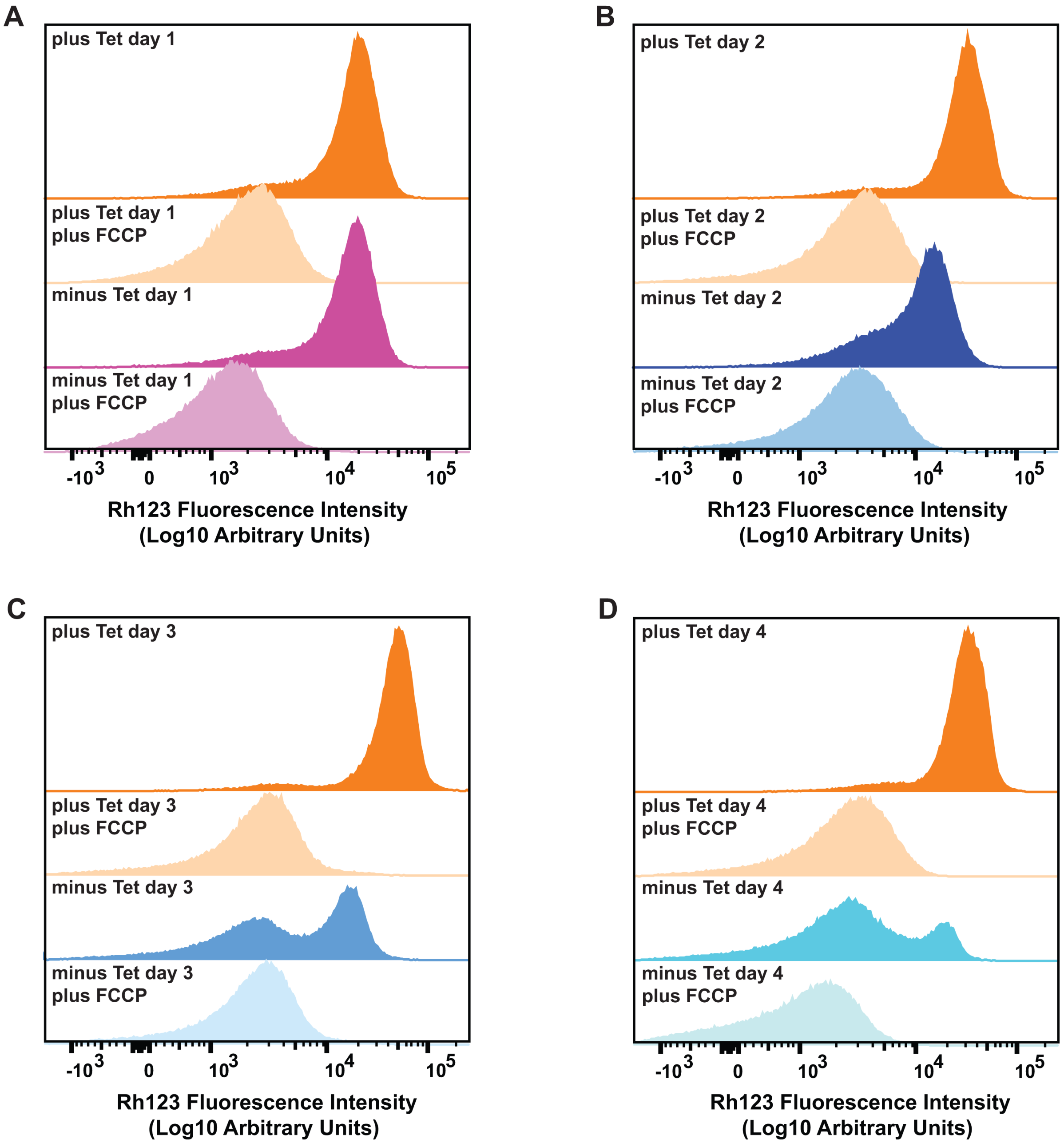

### Supplementary Figure 5

A

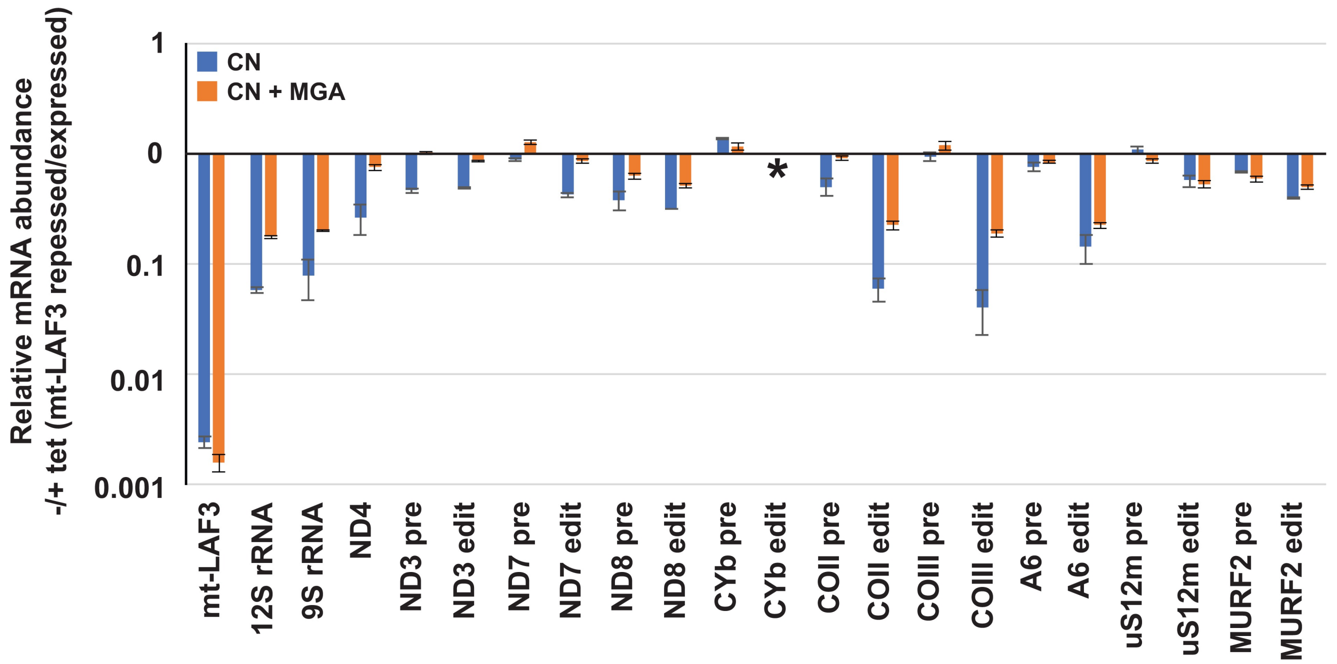


B

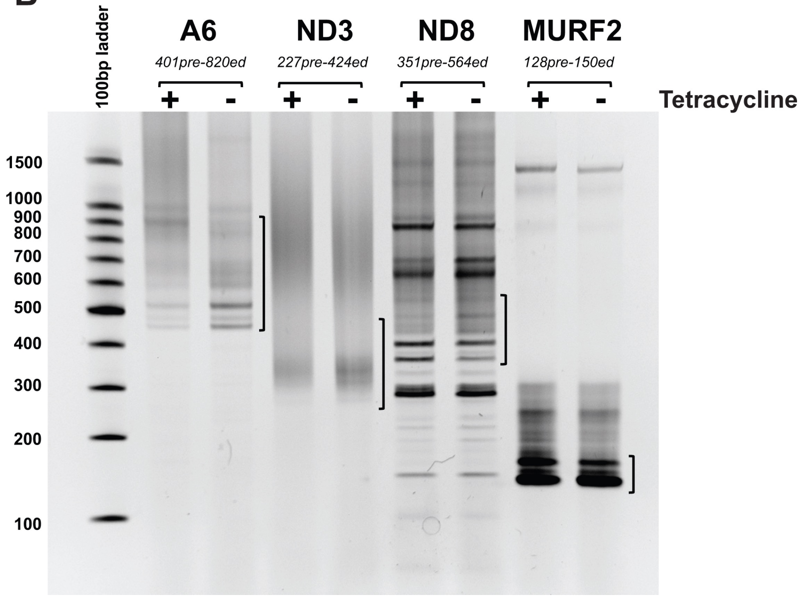
