## Supplementary Figure Legends and Tables for "mt-LAF3 is a pseudouridine synthase ortholog required for mitochondrial rRNA and mRNA gene expression in *Trypanosoma brucei*"

**Supplementary Figure 1. (A)** The presence of Pus family enzymes from each of the major eukaryotic families was assessed in an evolutionary tree for a diverse set of organisms built using several different markers (Maran et al., 2021). In general, Trypanosomatids have a similar profile of enzyme numbers per family when compared to the other organisms. **(B)** Amino acid identities and similarities between mt-LAF3 orthologs from Trypanosomatids, human, yeast, and *E.coli*.

**Supplementary Figure 2.** Clustal Omega alignment of *T. brucei* Tb927.9.3350/mt-LAF3 with related paralogs and orthologs from the RluA Pus family (Supplementary Table 3). The black box indicates the catalytic region, including the conserved catalytic aspartate (mt-LAF3 D211). An arginine residue thought to be key for RluA function (Hoang et al., 2006) is not conserved in mt-LAF3 (mt-LAF3 N209). The red box indicates an I/L residue found in most RluA family members that is also not conserved in mt-LAF3 and its direct Trypanosomatid orthologs (mt-LAF3 E316). Structural studies of mitochondrial ribosome assembly intermediates showed that this glutamate in mt-LAF3 forms hydrogen bonds with 12S rRNA that tilt the rRNA away from catalytic residues (Jaskolowski et al., 2020).

**Supplementary Figure 3.** Tet-regulated mt-LAF3-TAP and constitutive WT or D211A-V5 allele expression and repression in BF mt-LAF3 CN, BF mt-LAF3 CN + WT-V5, BF mt-LAF3 CN + D211A-V5, and BF mt-LAF3 CN + MGA cells were confirmed via Western blotting, probing with PAP or anti-V5 antibodies respectively, or for tubulin as a loading control (2 x 10^6^ cell equivalents per lane).

**Supplementary Figure 4.** The ΔΨm of BF mtLAF-3 CN cells was measured for four days following mt-LAF3 loss via tet withdrawal. ΔΨm was measured using rhodamine 123 (Rh123) and flow cytometry. Cells grown in the presence of tet, and thus expressing mt-LAF3, were included every day as a control. Cells were also pre-incubated with the protonophore trifluorocarbonylcyanide phenylhydrazone (FCCP) to define ΔΨm positive and negative cell populations. Representative analyses are shown, n = 2.

**Supplementary Figure 5.** RT-PCR analyses showing the impact of mt-LAF3 loss on RNA editing *in vivo*. **(A)** RT-qPCR assay for a range of mitochondrial rRNAs and mRNAs in BF mt-LAF3 CN and BF mt-LAF3 CN + MGA cells grown for 36 hours in the presence and absence of tet (both A and B datasets also shown in heatmap form in Figure 3). For each target amplicon, the relative change in RNA abundance was determined by using telomerase reverse transcriptase (TERT) mRNA as an internal control, with cell lines that have either mt-LAF3 repressed compared to the same cell line in which mt-LAF3 was expressed. Asterisk indicates that edited CYb was not detected in the CN cells grown either in the absence or presence of tet and we were therefore unable to quantify changes in its abundance. Data are shown as means ± SEM from five independent experiments. **(B)** RT-PCR products corresponding to full length pre-, partially- and fully- edited A6, ND3, ND8, and MURF2 mRNAs from BF CN cells in which mt-LAF3 is expressed (+) or repressed (-) in the presence or absence of tet. Brackets indicate the expected size ranges from pre-edited to fully edited amplicons as noted above for each mRNA.

**Supplementary Tables**

**Supplementary Table 1.** Sequences of oligonucleotides used in this study.

| **Primer number** | **Primer Description** | **Primer sequence** | **Reference** |
| --- | --- | --- | --- |
| 5355 | FOR oligo anneals 5’ of the β-tubulin locus integration site | GTACGCTGCTCATCTCGAAGCT | (McDermott et al., 2015) |
| 5356 | REV oligo anneals 3’ of ORF integrated into the β-tubulin locus | TTGGCCACACAACCCGGTGTTA | (McDermott et al., 2015) |
| 6391 | V5-tag in pHD1344tub (PAC)GW-Cterm3V5  REV sequencing primer | TCTAGATCTCGTGCTATCAAGACCGAGGA | (McDermott et al., 2015) |
| 9571 | GPEET splice-acceptor site in pHD1344tub (PAC)GW-Cterm3V5  FOR sequencing primer | GCTGCACGCGCCTTCGAGT | (McDermott et al., 2015) |
|  | Tb927.9.3350 knockout 5’ fragment forward | TCAGATTACATGAGAGCC | This study |
|  | Tb927.9.3350 knockout 5’ fragment forward nested | ATGGAGACCATGAAGGTC | This study |
|  | Tb927.9.3350 knockout 5’ fragment reverse | ATACGAAGTTATAAGCTTATCGTAGTCTTCTATAATACAAACAC | This study |
|  | Tb927.9.3350 knockout outer 3’ fragment forward | TTATGGATCCTAAGTGGGTCCGTGTTTTCACTTCTTTAC | This study |
|  | Tb927.9.3350 knockout outer 3’ fragment reverse | CTCACTTCCAACAGCAAT | This study |
|  | Tb927.9.3350 knockout outer 3’ fragment reverse nested | GAATCGCTCGCTACATCC | This study |
|  | Tb927.9.3350 knockout inner 3’ fragment forward | TTATGGATCCTAAGTGGGTCTATCAATGAATCCCAACG | This study |
|  | Tb927.9.3350 knockout inner 3’ fragment reverse | ATCTTCATTCTAGCAGCG | This study |
|  | Tb927.9.3350 gateway regulatable copy forward | GGGGACAAGTTTGTACAAAAAAGCAGGCTCCAAA ATGTTGCATGCCGTTCGC | This study |
|  | Tb927.9.3350 gateway regulatable copy reverse | GGGGACCACTTTGTACAAGAAAGCTGGGTC GCCCGGTATCAGCGGGT | This study |
|  | Tb927.9.3350 D211A site-directed mutagenesis | TTGTTTGTCATAACCTAGCCAGGGAAACTAGTGGGTG | This Study |
|  | Tb927.9.3350 D211A site-directed mutagenesis | CACCCACTAGTTTCCCTGGCTAGGTTATGACAAACAA | This Study |
|  | Tb927.9.3350 ORF BioMark forward | GTCAGCTACTGCACGAGAAA | This Study |
|  | Tb927.9.3350 ORF BioMark  reverse | GTCTGTCGGCATCATAGGTAAA | This Study |
| 4935 | MURF2 BioMark pre-edited forward | GATTTTAAGATTGGCTTTGATTGA | (Carnes et al., 2005) |
| 4934 | MURF2 BioMark pre-edited reverse | AATATAAAATCTAGATCAAACCATCACA | (Carnes et al., 2005) |
| 4969 | RPS12 BioMark pre-edited forward | CGACGGAGAGCTTCTTTTGAATA | (Carnes et al., 2005) |
| 4970 | RPS12 BioMark pre-edited reverse | CCCCCCACCCAAATCTTT | (Carnes et al., 2005) |
| 4928 | A6 BioMark pre-edited forward | TTGCCTTTGCCAAACTTTTAGAAG | (Carnes et al., 2005) |
| 4929 | A6 BioMark pre-edited reverse | ATTCTATAACTCCAAAATCACAACTTTCC | (Carnes et al., 2005) |
| 4965 | ND3 BioMark pre-edited forward | GAATGGGAGATGGGTTTTGG | (Carnes et al., 2005) |
| 4966 | ND3 BioMark pre-edited reverse | AACAAATCTCTTTACCCCCTTCAG | (Carnes et al., 2005) |
| 4942 | ND7 BioMark pre-edited forward | GCGGGCGGAGCATTATT | (Carnes et al., 2005) |
| 4943 | ND7 BioMark pre-edited reverse | GATCTACGGTCCCCTCTTTCCT | (Carnes et al., 2005) |
| 4953 | ND8 BioMark pre-edited forward | GATTTTTGCCAACGCATTCAG | (Carnes et al., 2018) |
| 4954 | ND8 BioMark pre-edited reverse | CACCGTGTAATTCTAAATTTCTCACTTC | (Carnes et al., 2018) |
| 4932 | CYb BioMark pre-edited forward | ATATAAAAGCGGAGAAAAAAGAAAG | (Carnes et al., 2005) |
| 4931 | CYb BioMark pre-edited reverse | CCCATATATTCTATATAAACAACCTGACA | (Carnes et al., 2005) |
| 4944 | COII BioMark pre-edited forward | ATTACAGTGTAACCATGTATTGACATT | (Carnes et al., 2005) |
| 4946 | COII BioMark pre-edited reverse | TTCATTACACCTACCAGGTTCTCT | (Carnes et al., 2005) |
| 4938 | COIII BioMark pre-edited forward | GAAACCAGATGAGATTGTTTGCA | (Carnes et al., 2005) |
| 4939 | COIII BioMark pre-edited reverse | TTCATTCCAACTAAACCCTTTCC | (Carnes et al., 2005) |
| 11040 | TERT BioMark forward | GAGCGTGTGACTTCCGAAGG | (Brenndorfer and Boshart, 2010) |
| 11041 | TERT BioMark reverse | AGGAACTGTCACGGAGTTTGC | (Brenndorfer and Boshart, 2010) |
| 4933 | MURF2 BioMark edited forward | GATTTTAATGTTTGGTTGTTTTAATTTAG | (Carnes et al., 2005) |
| 4934 | MURF2 BioMark edited reverse | AATATAAAATCTAGATCAAACCATCACA | (Carnes et al., 2005) |
| 4967 | RPS12 BioMark edited forward | CGTATGTGATTTTTGTATGGTTGTTG | (Carnes et al., 2005) |
| 4968 | RPS12 BioMark edited reverse | ACACGTCGGTTACCGGAACT | (Carnes et al., 2005) |
| 5071 | A6 BioMark edited forward | GATTTATTTTGGTTGCGTTTGTTATTATG | (Carnes et al., 2005) |
| 5072 | A6 BioMark edited reverse | CAAACCAACAAACAAATACAAATCAAAC | (Carnes et al., 2005) |
| 4965 | ND3 BioMark edited forward | GAATGGGAGATGGGTTTTGG | (Carnes et al., 2005) |
| 4966 | ND3 BioMark edited reverse | AACAAATCTCTTTACCCCCTTCAG | (Carnes et al., 2005) |
| 4942 | ND7 BioMark edited forward | GCGGGCGGAGCATTATT | (Carnes et al., 2005) |
| 4943 | ND7 BioMark edited reverse | GATCTACGGTCCCCTCTTTCCT | (Carnes et al., 2005) |
| 6402 | ND8 BioMark edited forward | AATTTGCCCTAGTTTAGCATTGG | (Carnes et al., 2018) |
| 6403 | ND8 BioMark edited reverse | ATCAATCCGCAAAACGATGA | (Carnes et al., 2018) |
| 4930 | CYb BioMark edited forward | AAATATGTTTCGTTGTAGATTTTTATTATTT | (Carnes et al., 2005) |
| 4931 | CYb BioMark edited reverse | CCCATATATTCTATATAAACAACCTGACA | (Carnes et al., 2005) |
| 4944 | COII BioMark edited forward | ATTACAGTGTAACCATGTATTGACATT | (Carnes et al., 2005) |
| 4945 | COII BioMark edited reverse | ATTTCATTACACCTACCAGGTATACAA | (Carnes et al., 2005) |
| 4936 | COIII BioMark edited forward | TTGTGTTTTATTACGTTGTATCCAGTATTG | (Carnes et al., 2005) |
| 4937 | COIII BioMark edited reverse | CGAAAGCAAACTCACAACACAAA | (Carnes et al., 2005) |
| 5067 | COI BioMark forward | CCCGATATGGTATTTCCTCGTATAAA | (Carnes et al., 2005) |
| 5068 | COI BioMark reverse | CCCCCATACCCTCTTCAGTCA | (Carnes et al., 2005) |
| 5069 | ND4 BioMark forward | CAATCTGACCATTCCATGTGTGA | (Carnes et al., 2005) |
| 5070 | ND4 BioMark reverse | TTTCAGCACAATACTTGCTAATAAAACA | (Carnes et al., 2005) |
| 10662 | 12S rRNA BioMark forward | GGGCAAGTCCTACTCTCCTTTACAAAG | (Aphasizheva et al., 2011) |
| 10663 | 12S rRNA BioMark reverse | TGAACAATCAATCATGGTAATAAGTAGACGATG | (Aphasizheva et al., 2011) |
| 10664 | 9S rRNA BioMark forward | ATTAGATTGTTTTGTTAATGCTATTAGATG | (Aphasizheva et al., 2011) |
| 10665 | 9S rRNA BioMark reverse | ACGGCTGGCATCCATTTC | (Aphasizheva et al., 2011) |
| 3623 | FOR ND8 pre-edited/edited RT-PCR | CAATTTAATAATTTTAAGTTTTGG | (Carnes et al., 2022) |
| 3624 | REV ND8 pre-edited/edited RT-PCR | TAGTCAAAATTTAATTTCACCGTG | (Carnes et al., 2022) |
| 7263 | FOR ND3 pre-edited/edited RT-PCR | CCTCGCCTTTTTACTTTAGTTTGTTATCA | (Carnes et al., 2022) |
| 4805 | REV ND3 pre-edited/edited RT-PCR | TAATATGTATAATACAAC | (Carnes et al., 2022) |
| 6204 | FOR MURF2 pre-edited/edited RT-PCR | ATAGAAAGGTATATAATCTATAATG | (Guo et al., 2010) |
| 4934 | REV MURF2 pre-edited/edited RT-PCR | AATATAAAATCTAGATCAAACCATCACA | (Carnes et al., 2005) |
| 3704 | FOR A6 pre-edited/edited RT-PCR | AAAAATAAGTATTTTGATATTATTAAAG | (Schnaufer et al., 2001; Guo et al., 2010) |
| 3580 | REV A6 pre-edited/edited RT-PCR | TATTATTAACTTATTTGATC | (Schnaufer et al., 2001; Guo et al., 2010) |

**Supplementary Table 2.** Antibodies used in this study.

| **Antibody name** | **Raised in** | **Dilution** | **Source** |
| --- | --- | --- | --- |
| V5 Epitope Tag Monoclonal Antibody | Mouse | 1/5000 WB | ThermoFisher Scientific  R960-25 |
| Anti-Mouse IgG (H+L)-HRP Conjugate | Goat | 1/5000 WB | Bio-Rad 1706516 |
| PAP Soluble Complex Antibody | Rabbit | 1/2000 WB | Sigma P1291 |
| Anti-Tubulin [YL1/2] Monoclonal Antibody-Loading Control | Rat | 1/5000 WB | Abcam ab6160 |
| Recombinant Protein G-HRP | N/A | 1/5000 WB | ThermoFisher Scientific  101223 |

**Supplementary Table 3.** Gene and protein IDs for sequences used in multi-species alignment of mt-LAF3 with its paralogs and orthologs.

| **Species** | **GeneID** | **Protein** | **Uniprot Protein ID** |
| --- | --- | --- | --- |
| *Trypanosoma brucei brucei* TREU927 | Tb927.9.3350 | mt-LAF3 | Not present in Uniprot database |
| *Trypanosoma cruzi* Dm28c | BCY84_12019 | mtLAF-3 ortholog | Not present in Uniprot database |
| *Leishmania major* Friedlin | LMJFC_010007900 | mtLAF-3 ortholog | Not present in Uniprot database |
| *Trypanosoma brucei brucei* TREU927 | Tb927.3.2130 | mtLAF-3 paralog | Q57Z98 |
| *Trypanosoma cruzi* Dm28c | BCY84_01007 | Tb927.3.2130 ortholog | Not present in Uniprot database |
| *Leishmania major* Friedlin | LMJFC_350064900 | Tb927.3.2130 ortholog | Not present in Uniprot database |
| *Saccharomyces cerevisiae* S288C | PUS5/YLR165C | PUS5 | Q06244 |
| *Homo sapiens* RPUSD3 | RPUSD3/HGNC:28437 | RPUSD3 | Q6P087 |
| *Homo sapiens* RPUSD4 | RPUSD4/HGNC:25898 | RPUSD4 | Q96CM3 |
| *E. coli* RluA strain K12 | rluA/EG12609 (EcoCyc) | RluA | P0AA37 |

**Supplementary References**

Aphasizheva, I., Maslov, D., Wang, X., Huang, L., Aphasizhev, R., 2011. Pentatricopeptide repeat proteins stimulate mRNA adenylation/uridylation to activate mitochondrial translation in trypanosomes. Mol Cell 42, 106-117. doi: 10.1016/j.molcel.2011.02.021

Brenndorfer, M., Boshart, M., 2010. Selection of reference genes for mRNA quantification in Trypanosoma brucei. Molecular and biochemical parasitology 172, 52-55. doi: 10.1016/j.molbiopara.2010.03.007

Carnes, J., Gendrin, C., McDermott, S.M., Stuart, K., 2022. KRGG1 function in RNA editing in Trypanosoma brucei. RNA. doi: 10.1261/rna.079418.122

Carnes, J., McDermott, S.M., Stuart, K., 2018. RNase III Domain of KREPB9 and KREPB10 Association with Editosomes in Trypanosoma brucei. mSphere 3. doi: 10.1128/mSphereDirect.00585-17

Carnes, J., Trotter, J.R., Ernst, N.L., Steinberg, A., Stuart, K., 2005. An essential RNase III insertion editing endonuclease in Trypanosoma brucei. Proc Natl Acad Sci U S A 102, 16614-16619. doi: 10.1073/pnas.0506133102

Guo, X., Ernst, N.L., Carnes, J., Stuart, K.D., 2010. The zinc-fingers of KREPA3 are essential for the complete editing of mitochondrial mRNAs in Trypanosoma brucei. PloS one 5, e8913. doi: 10.1371/journal.pone.0008913

Hoang, C., Chen, J., Vizthum, C.A., Kandel, J.M., Hamilton, C.S., Mueller, E.G., Ferre-D'Amare, A.R., 2006. Crystal structure of pseudouridine synthase RluA: indirect sequence readout through protein-induced RNA structure. Mol Cell 24, 535-545. doi: 10.1016/j.molcel.2006.09.017

Jaskolowski, M., Ramrath, D.J.F., Bieri, P., Niemann, M., Mattei, S., Calderaro, S., Leibundgut, M., Horn, E.K., Boehringer, D., Schneider, A., Ban, N., 2020. Structural Insights into the Mechanism of Mitoribosomal Large Subunit Biogenesis. Mol Cell 79, 629-644 e624. doi: 10.1016/j.molcel.2020.06.030

Maran, S.R., de Lemos Padilha Pitta, J.L., Dos Santos Vasconcelos, C.R., McDermott, S.M., Rezende, A.M., Silvio Moretti, N., 2021. Epitranscriptome machinery in Trypanosomatids: New players on the table? Mol Microbiol 115, 942-958. doi: 10.1111/mmi.14688

McDermott, S.M., Carnes, J., Stuart, K., 2015. Identification by Random Mutagenesis of Functional Domains in KREPB5 That Differentially Affect RNA Editing between Life Cycle Stages of *Trypanosoma brucei*. Molecular and cellular biology 35, 3945-3961. doi: 10.1128/MCB.00790-15

Schnaufer, A., Panigrahi, A.K., Panicucci, B., Igo, R.P., Jr., Wirtz, E., Salavati, R., Stuart, K., 2001. An RNA ligase essential for RNA editing and survival of the bloodstream form of Trypanosoma brucei. Science 291, 2159-2162. doi: 10.1126/science.1058955
